# Unbiased Long-Read Whole-Genome Sequencing Enables High-Resolution Mapping of Transgene Concatenation and Off-target Genomic Disruption in a Mouse Model

**DOI:** 10.64898/2026.05.15.725597

**Authors:** Maanik Mehta, Kashif Ahmed, Rowaida Hussein, Erika Tavares, Zorana Berberovic, Rabiat Adele, Abigail D’Souza, Bin Gu, Michael D. Wilson, Evgueni Ivakine, Philippe P. Monnier, Elise Héon, Ajoy Vincent

**Author notes:** Corresponding Author: Dr. Ajoy Vincent.

## Abstract

Transgenic mouse models are indispensable for dissecting disease mechanisms, yet their interpretability is frequently compromised by cryptic genomic alterations introduced during transgenesis. Thus, robust quality control strategies are needed to elucidate integration architecture and evaluate model performance when such unintended events occur. Here, we applied unbiased whole-genome long-read sequencing using the PacBio Revio to investigate a mouse model exhibiting unexpected transgene silencing, originally designed to recapitulate autosomal-dominant hereditary macular dystrophy driven by upregulation of a *ZZEF1-ALOX15* fusion gene. Long-read sequencing analysis revealed a ≥29 kb head-to-tail concatemer containing at least three transgene-derived copies. Reconstruction of transgene-genome junctions revealed unintended off-target integration of the concatemer into the calcium-sensing receptor gene (*Casr*), along with exogenous *E. coli* DNA, that together defined final transgene architecture. Aggregate 5-methylcytosine profiling identified hypermethylation of the transgene promoter across the concatemer and additional phenotyping suggested disruption of endogenous CaSR function resulting from the rearrangement. Our workflow enabled direct detection of transgene concatenation and off-target mapping. These findings highlight the power of whole-genome LRS in uncovering hidden genomic complexity, resolving aberrant phenotypes and enhancing the reliability of *in vivo* disease modelling, supporting its incorporation as a potentially scalable quality control tool for genetically engineered animal models.

## Introduction

The development of genetically engineered mice (GEMs) over the past decade has heralded our unprecedented ability to interrogate the complex mechanisms of human disease. Today, a suite of powerful editing tools can enable precise, “on-target”, manipulations of the murine genome, ranging from single base-pair modifications to multi-kilobase insertions, deletions and duplications^1–4^. While the CRISPR-Cas9 system remains the preferred gene-editing strategy, its efficiency in facilitating large on-target insertions is limited^5^. Moreover, unintended rearrangements between transgenes and the host genomes are frequently observed during transgenesis and can confound phenotypic characterization by disrupting endogenous gene function^6–9^. These include the concatenation of transgenic DNA during cell microinjection, whereby copies of plasmid recombine into tandem arrays prior to genomic integration^10,11^.

Identification of concatemers is critical to deciphering model viability, as their formation can result in aberrant transgene copy number, loss of plasmid elements, or even repeat-induced gene silencing (RIGS)^10,12,13^. Nonetheless, these unintended outcomes may be missed by traditional PCR-based methods due to technical limitations in amplifying large and/or repetitive transgenes, the loss of primer binding sites, multiple integration sites, or preservation of integration junctions^11,14–16^. Conventional short-read next-generation sequencing technologies, which can generally map integration sites through split/discordant read analysis, similarly struggle to characterize repetitive transgene rearrangements when these structures exceed the read length yields of 150 bp that most platforms provide, even with sequencing coverage as high as 250×^17–20^. Thus, robust quality control measures that permit reliable and efficient detection of such effects are needed.

In recent years, long-read sequencing (LRS) platforms such as Pacific Biosciences (PacBio) or Oxford Nanopore Technologies (ONT) have emerged as novel methods to elucidate transgene architecture. The primary benefit of LRS is its ability to generate median read lengths of 10-100 kb, capturing repetitive DNA and concatemer-host genome junctions within a single read^21,22^. An added advantage for both PacBio and ONT over short-read sequencing is their ability to perform onboard methylation profiling, concurrently informing transgene structure and its expression potential^23–25^. Early transgene mapping attempts using whole-genome LRS yielded only a minute fraction of reads capturing the transgene-genome junctions, demonstrating considerable inefficiencies with these sequencing workflows^26,27^. Accordingly, LRS is frequently combined with Cas9-based enrichment strategies, which selectively cleave DNA within the transgene or around a predetermined locus for adaptor ligation and sequencing^28^. While these techniques achieve exceptional coverage of the integration locus, their targeted nature offers limited utility in examining unwanted editing outcomes that are not physically linked to the captured transgene sequence^9,29,30^. Thus, when the underlying mechanism of model failure or inefficiency is unknown, a comprehensive genome-wide strategy offering non-targeted resolution would be particularly advantageous^31^.

Here, we describe the application of unbiased whole-genome LRS to investigate transgene silencing in a mouse model, designed to recapitulate a form of autosomal-dominant hereditary macular dystrophy (HMD) driven by the ectopic overexpression of a 3.3 kb *ZZEF1-ALOX15* fusion gene^32^. Previous *in vivo* electroporation of this *ZZEF1-ALOX15* chimera demonstrated retinal photoreceptor damage, motivating the generation of a stable mouse line for mechanistic studies^32^. Thus, a Cre-inducible expression vector (*pCAG-ZZEF1-ALOX15-T2A-L2T*) containing the chimera was targeted to the *ROSA26* safe harbor locus in mice using CRISPR-Cas9. The generated line was subsequently crossed to retinal-specific Cre-driver strains. Despite passing standard quality control at the generation facility, the resulting mice displayed no evidence of transgene activation, prompting deeper investigation. We performed PacBio^®^ high-fidelity (HiFi) sequencing of the mouse model, identifying a complex multi-copy transgene concatemer with unintended off-target integration into the calcium-sensing receptor gene (*Casr)* and hypermethylation of the transgene promoter in ear-notch DNA. Systemic evaluation revealed altered calcium homeostasis in transgenic mice as a functional consequence of the rearrangement. Together, these findings provide a mechanistic framework for the failure of transgene activation in this model and demonstrate the broad utility of LRS as a quality control tool for GEM characterization.

## Materials and Methods

### Mouse Housing and Maintenance

Mice were housed at The Centre for Phenogenomics (TCP, Toronto, Canada) in accordance with the Canadian Council on Animal Care (CCAC) guidelines and mandates established by Ontario’s Animals for Research Act. Animals were provided *ad libitum* access to water and standard chow and were maintained under aseptic conditions with a 12-hour light/dark cycle. All animal protocols were approved by the TCP Animal Care Committee (AUP 29-0368).

### Generation of Transgenic Mouse Model

A 19 kb construct (*pCAG-ZZEF1-ALOX15-T2A-L2T*) containing the 3.3 kb *ZZEF1-ALOX15* fusion gene and a Luciferase-tdTomato (L2T) fusion reporter, under control of the CMV early enhancer/chicken β-actin (CAG) promoter was generated using InFusion cloning (Figure 1A). A loxP-flanked stop cassette (LSL) immediately downstream of the CAG promoter was used to enable Cre-inducible activation of the transgene. The backbone plasmid contained an ampicillin resistance cassette as well as 5′ and 3′ homology arms (800-bp and 4.3 kb, respectively) to direct integration of the construct into the *ROSA26* locus. Of the 19 kb construct, the cassette for genomic integration (excluding backbone elements and homology arms) spanned 11 kb. The final construct (Data File S1) was validated by Sanger sequencing at The Centre for Applied Genomics (TCAG, The Hospital for Sick Children, Toronto, Canada). Construct functionality was assessed by co-transfection with pCMV-Cre (Addgene plasmid #123133) in HEK293T cells using Lipofectamine 3000 (L3000001, Invitrogen, Carlsbad, California, USA), followed by imaging to detect tdTomato fluorescence 24 hours post-transfection.

**Figure 1.**
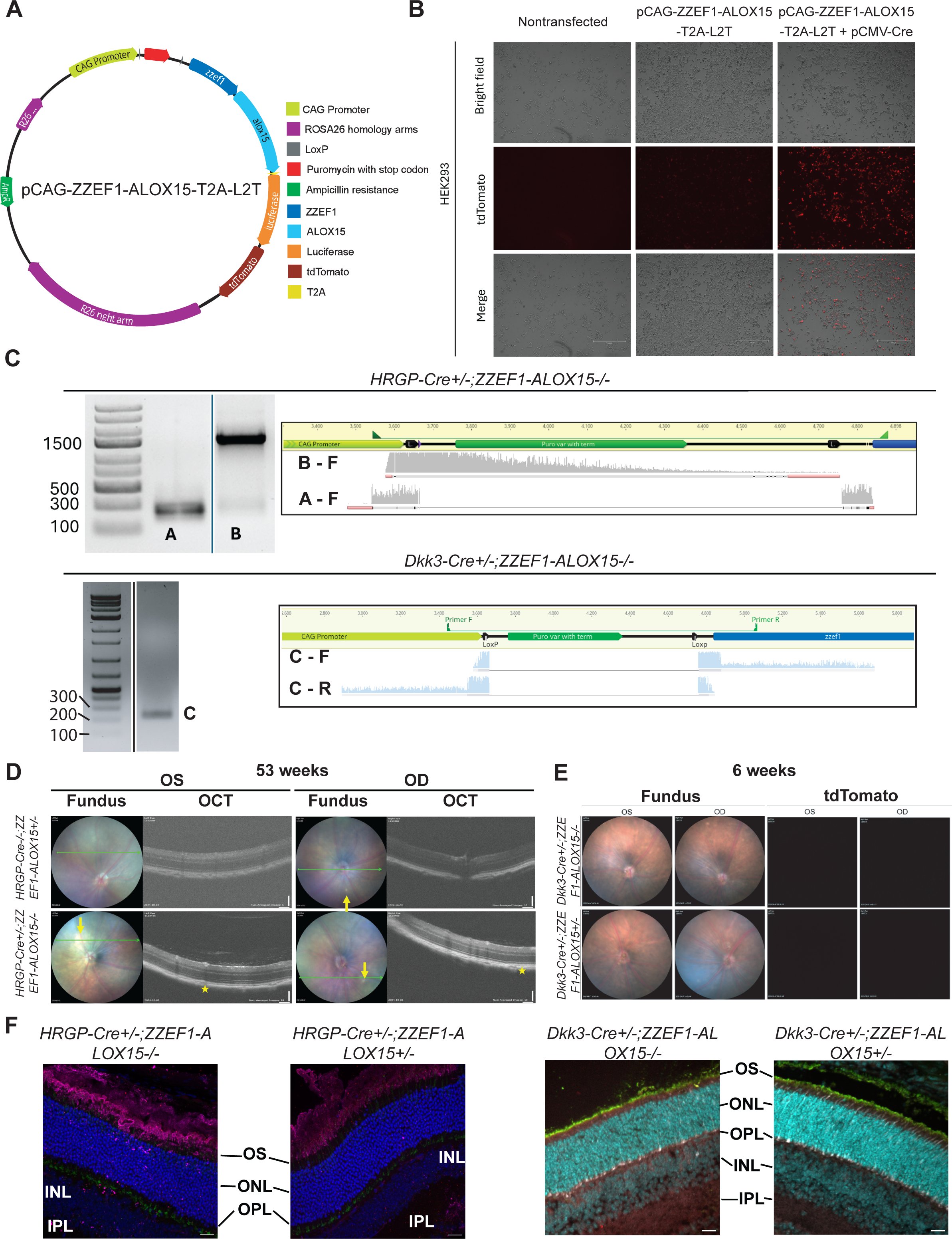
OS (panels D/E), left eye; OD, right eye; OCT, optical coherence tomography; OS (panel F), outer segment; ONL, outer nuclear layer; OPL, outer plexiform layer; INL, inner nuclear layer; IPL, inner plexiform layer. **(A)** Schematic of pCAG-ZZEF1-ALOX15-T2A-L2T designed to express the *ZZEF1-ALOX15* fusion gene, luciferase and tdTomato under the control of the CAG promoter. **(B)** In vitro validation of the construct. Co-transfection of HEK293T cells with pCMV-Cre activated tdTomato reporter expression, confirming construct functionality. Images were captured at 10× original magnification; scale bars, 750 μm. **(C)** Gel electrophoresis and Sanger sequencing alignment (Geneious Prime) following PCR genotyping of retinal DNA from *HRGP-Cre^+/-^ ;ZZEF1-ALOX15^+/-^* and *Dkk3-Cre^+/-^;ZZEF1-ALOX15^+/-^* mice, showing excision of the loxP-flanked stop cassette. **(D)** Top panel: Fundus and OCT examination of both eyes of an *HRGP-Cre^+/-^;ZZEF1-ALOX15^-/-^*littermate control at week 54. The small yellow spot in the right eye fundus is suggestive of drusenoid lesion development. Bottom panel: Both eyes of an *HRGP-Cre^+/-^;ZZEF1-ALOX15^+/-^* mouse showing dots on fundus corresponding to hyper-reflective drusenoid lesions (*) at the RPE. Retinal lamination appears preserved across both transgenic and control mice. **(E)** Fundus imaging of both eyes at week 5 showing a littermate control (*Dkk3-Cre^+/-^;ZZEF1-ALOX15^-/-^*) in the top panel and a transgenic mouse (*Dkk3-Cre^+/-^;ZZEF1-ALOX15^+/-^*) in the bottom panel. Both eyes appear unremarkable on examination. Imaging with a tdTomato excitation filter over the same region reveals no detectable fluorescence. **(F)** Cryo-immunohistochemistry (cryo-IHC) showing absence of tdTomato reporter in both transgenic lines. In *HRGP-Cre^+/-^;ZZEF1–ALOX15^+/-^* sections (8 weeks), nuclei were stained with DAPI (cyan), rods with rhodopsin (purple), cones with cone arrestin (green), and tdTomato (red). In *Dkk3-Cre^+/-^;ZZEF1–ALOX15^+/-^* sections (11 days), nuclei were stained with DAPI (cyan), rods with rhodopsin (green), cones with cone arrestin (white), and tdTomato (red). Images were captured at 40× original magnification; scale bars, 20 μm.

An editing mix containing recombinant Cas9 protein, a guide RNA targeting the *ROSA26* locus (Supplementary Table S1), and the donor plasmid was injected into C57BL/6J embryos at the 2-cell stage at the McGill Integrated Core for Animal Modeling (MICAM, McGill University, Montreal, Canada).

Resulting B6.Gt(ROSA)26Sor^tm^^1(CAG-ZZEF^^1/ALOX^^15)1Vin^ founder mice, referred to as *ZZEF1-ALOX15* mice hereinafter, underwent routine PCR-based quality control by MICAM, using primers spanning the 5′ and 3′ junctions to verify construct integration (Supplementary Table S2). The founder line that passed this validation was then rederived at the TCP and bred to establish stable lines. PCR genotyping of the tdTomato reporter confirmed successful germline transmission (Supplementary Table S2). Offspring from this line were then crossed by our group to the *HRGP-Cre* driver strain^33^, obtained from the Jackson Laboratory (Bar Harbor, USA; strain #032911), to selectively induce transgene activation in the cone-photoreceptor cell population.

To enhance the probability of detecting transgene activation, transgenic mice were also crossed to the *Dkk3-Cre* line^34^ (generated by Dr. Takahisa Furukawa, University of Osaka, Osaka, Japan), which mediates expression in almost all retinal progenitor cells.

### Retinal Phenotyping of Cre-Activated Litters

Retinae from transgenic mice carrying the *ZZEF1-ALOX15* construct and either the cone-photoreceptor specific *HRGP-Cre (HRGP-Cre^+/-^;ZZEF1-ALOX15^+/-^*) or the pan-retinal *Dkk3-Cre* (*Dkk3-Cre^+/-^;ZZEF1-ALOX15^+/-^*) driver alleles were harvested at week 28 and 77, respectively for DNA extraction. PCR from these extracts was performed using AmpliTaq Gold™ 360 Master Mix (Applied Biosystems; 4398881) to verify Cre-mediated excision of the LSL cassette in our construct (Supplementary Table S2). Bands corresponding to the expected amplicon size were gel-extracted and re-amplified prior to Sanger sequencing.

Transgenic mice (*HRGP-Cre^+/-^;ZZEF1-ALOX15^+/-^*and *Dkk3-Cre^+/-^;ZZEF1-ALOX15^+/-^*) underwent descriptive ocular phenotyping alongside littermate controls. During *in vivo* imaging, mice were anesthetized using ketamine (100 mg/kg) and xylazine (10 mg/kg), which were administered intraperitoneally (0.10 mL/10 g body weight). Assessments included fundus photography and optical coherence tomography (OCT) using the Phoenix MICRON 5 camera (Phoenix Research Laboratories, Pleasanton, CA) at months 1, 2, 3, 6, 9 and 12 for the *HRGP-Cre* x *ZZEF1-ALOX15* cohort [*HRGP-Cre^+/-^;ZZEF1-ALOX15^+/-^*(n = 4); *HRGP-Cre^+/-^*;*ZZEF1-ALOX15^-/-^* (n = 2)]. OCT examination provides ultrastructural imaging of the retina, allowing detection of morphological changes to the retinal layers as those previously observed in human subjects^32^. Additionally, live fundus imaging with a tdTomato excitation/emission filter set (Phoenix MICRON 5) was performed on the *Dkk3-Cre* cohort at week 5 (n = 1, *Dkk3-Cre^+/-^ ;ZZEF1-ALOX15^+/-^*; n = 1, *Dkk3-Cre^+/-^;ZZEF1-ALOX15^-/-^*). For *ex vivo* evaluation, the retina and retinal pigment epithelium (RPE) tissue was collected from a transgenic mouse and control littermate from each line (*Dkk3-Cre^+/-^;ZZEF1-ALOX15^+/-^* and *Dkk3-Cre^+/-^;ZZEF1-ALOX15^-/-^* at P11; *HRGP-Cre^+/-^;ZZEF1-ALOX15^+/-^* and *HRGP-Cre^-/-^;ZZEF1-ALOX15^+/-^*at week 8) for immunohistochemistry (IHC) to assess reporter expression and cellular morphology as previously described^32^ except that retinal cryosections were incubated with tdTomato antibodies (Origene; AB8181-200, 1:400) instead of GFP in combination with Cone Arrestin (Millipore-Sigma; AB15282; 1:1000), and Rhodopsin (Sigma-Aldrich; O4886, 1:10,000)^32^. All comparisons were made between transgenic mice and littermate controls lacking either *Cre* or transgene.

In addition to ocular phenotyping, total RNA was isolated from the retinae of *HRGP-Cre^+/-^ ;ZZEF1-ALOX15^+/-^*and *Dkk3-Cre^+/-^;ZZEF1-ALOX15^+/-^* mice (weeks 152 and 77, respectively) using the Monarch Total RNA Miniprep Kit (New England Biolabs) as per the manufacturer’s protocol. Eluates were treated with DNase I (Thermo Scientific) for 30 minutes at 37°C and subsequently purified using the PuroSPIN Total RNA Purification Kit (Luna Nanotech). The resulting RNA was reverse transcribed to cDNA using the LunaScript RT SuperMix kit (New England Biolabs), with 100 ng of RNA in a 20 µL reaction. To assess transgene expression at the RNA level, PCR targeting the *ZZEF1-ALOX15* fusion junction and Cre recombinase (Supplementary Table S2) was performed with 2× Taq FroggaMix (FroggaBio) using 5 ng of resulting cDNA from each sample. PCR products were resolved on a 2.0% agarose gel and imaged using the ChemiDoc Imaging system (Bio-Rad, Hercules, CA, USA). Bands of interest were excised, isolated by centrifugation (10,000 RCF for 10 minutes) and Sanger sequenced at TCAG.

### Whole-genome LRS Analysis of *ZZEF1-ALOX15* Transgenic Mice

Genomic DNA extracted from an ear notch of a *ZZEF1-ALOX15^+/-^* transgenic mouse (F_4_ descendant of the original founder) was sequenced using the PacBio Revio at TCAG. Raw barcode-assigned HiFi reads provided in a BAM file were then mapped and analyzed for initial assessment. Reads were converted to FASTQ format using SAMtools v1.21^35^ and aligned to the mouse reference genome (GRCm39) using minimap2 v2.24^36^. We used the “map-pb” preset in minimap2 with soft-clipping enabled to retain sequences failing to map to the reference genome, ensuring that plasmid-derived sequences were not eliminated. Additional quality filtering was not applied after alignment to preserve exogenous reads in the event of significant transgene rearrangement. The resulting BAM file was then indexed with SAMtools and analyzed using the Integrative Genomics Viewer (IGV) v2.19.6^37^. To determine average genome-wide coverage and the number of mapped reads, mapping statistics were generated using the SAMtools “stats” command. Coverage was calculated by dividing the total bases mapped among barcode-assigned and primary mapped reads (CIGAR) by the haploid GRCm39 genome size.

To analyze transgene structure, reads containing transgenic DNA were identified and extracted by aligning raw reads to a multi-FASTA file (Data File S2) containing independent “bait” sequences of vector components (CAG promoter, *ZZEF1*, *ALOX15*, LSL cassette, tdTomato, luciferase, and backbone plasmid elements) using minimap2 with the “map-pb” preset and soft clipping enabled. Reads extracted by more than one vector component were deduplicated by read ID. To avoid capturing endogenous orthologs, reads identified by *ZZEF1* or *ALOX15* baits were only retained if they were supported by at least one additional vector component. Resulting reads were then imported into Geneious Prime v2025.1.3 (www.geneious.com). This baiting strategy was applied separately to the barcode-assigned and -unassigned HiFi read datasets. All baited reads were manually inspected in Geneious Prime for transgene elements before downstream analysis. Here, we employed the “Annotate and Predict” tool to resolve transgene-associated elements within individual reads against a database of manually curated plasmid features, with a ≥ 50% similarity threshold applied to identify vector components that may have been corrupted during integration. To perform a preliminary assessment of the *ROSA26* locus, transgene-derived reads identified by this strategy were removed from the initial alignment before re-visualizing coverage across the region on IGV.

For determination of the integration site, transgene-derived reads with terminal sequences that did not match exogenous DNA were queried against the mouse genome using BLAT to identify 5′ and 3′ junctions. Any unresolved sequences that remained within these reads were further annotated using BLAST^38^. Annotated junctional reads were aligned to the candidate integration site using the minimap2 plugin within Geneious Prime. A final integration site map with partial assembly of the concatemer was then generated using a consensus sequence of mapped reads. Junction sequences were aligned with the corresponding host-genome and exogenous reference sequences to inspect repair signatures including homologies/microhomologies, local rearrangements, insertions/templated insertions at each integration breakpoint. *De novo* assembly of the entire transgene-derived dataset was attempted using Flye v2.9.6^39^ and the native Geneious Prime *de novo* assembler. *In silico* analysis of potential genome-wide off-target Cas9 editing sites was performed using Cas-OFFinder^40^ (≤ 4 mismatches). Concurrently, genome-wide structural variants (SVs) were called using pbsv v2.6.2 (https://github.com/PacificBiosciences/pbsv) across the GRCm39 reference genome, with variants passing the standard pbsv quality filter (FILTER=PASS) retained for downstream analysis. Potential editing sites were then cross-referenced with the retained SV calls to explore potential Cas9-mediated rearrangements.

To assess site-level CpG methylation across the CAG promoter, raw HiFi reads identified by the CAG sequence bait were extracted using SAMtools. Resulting reads were subsequently aligned to the promoter reference and indexed using pbmm2 v26.1.0. Read-level 5-methylcytosine (5mC) calls generated by Jasmine (v2.3.0) were then parsed using pb-CpG-tools v3.0.0 (https://github.com/PacificBiosciences/pb-CpG-tools/), where CpG dinucleotides with modification scores ≥50 were classified as methylated.

Key tools, commands, and parameters utilized for the whole genome LRS analysis are provided in Data File S3.

### Validation of Off-Target Transgene Integration

Confirmatory PCR spanning the 5′ and 3′ integration junctions was performed with 2× Taq FroggaMix (FroggaBio) to verify allele status in all transgenic and control mice undergoing biochemical and *Casr* expression analysis (Supplementary Table S2).

*Casr* expression in the retina was assessed using reverse transcription quantitative polymerase chain reaction (RT-qPCR). RNA was isolated from both retinae of a female *ZZEF1-ALOX15* mouse (*Dkk3-Cre^-/-^;ZZEF1-ALOX15^+/-^*, week 77) and a related sex-matched control (maternal first-cousin; *Dkk3-Cre^-/-^;ZZEF1-ALOX15^-/-^*, week 74). RNA from the control mouse was isolated using the RNeasy Plus Mini Kit (QIAGEN), following homogenization through a QIAshredder column according to the manufacturer’s protocol. RNA from the *Dkk3-Cre^-/-^;ZZEF1-ALOX15^+/-^* mouse was isolated as described earlier (refer to *Retinal Phenotyping of Cre-Activated Litters* section). Resulting RNA samples were reverse transcribed using the LunaScript RT SuperMix Kit (New England Biolabs). To explore expression differences in the transgenic mouse relative to wild-type control, qPCR targeting exons 2-3 of *Casr* was performed using 5 ng of resulting cDNA, with each sample run in technical triplicates and normalized to *Ppib* (Supplementary Table S2).

Whole blood from the saphenous vein and spot urine were collected from transgenic mice (*ZZEF1-ALOX15^+/-^,* n = 9 for both blood and urine) and controls lacking the transgene (*ZZEF1-ALOX15^-/-^,* n = 10 for blood and n = 8 for urine, as two aged mice failed to produce urine) for calcium analysis. Animals included males and females with ages ranging from 23-111 weeks and 30-122 weeks for serum and urine analysis, respectively. Whole blood was kept at 4°C prior to centrifugation at 1300 RCF for 15 minutes to isolate serum, of which a minimum of 15 μL was subsequently frozen at -80°C until testing. A minimum of 25 μL of urine was stored at -80°C until time of testing. Isolated serum calcium and urinary calcium/creatinine concentrations were measured using the Beckman Coulter AU480 chemistry analyzer. Boxplots and statistical analyses were generated using R v4.5.1^41^ in RStudio v2025.05.1.513^42^ with tidyverse v2.0.0^43^, ggplot2 v3.5.2^44^ and ggpubr v0.6.1^45^. Associations between genotype and serum calcium or urinary calcium-to-creatinine ratios were analyzed using multiple linear regression models adjusted for age and sex, with genotype effect reported with 95% confidence intervals.

## Results

### Functional Assays of *ZZEF1-ALOX15* Mice Suggest Transgene Silencing

Co-transfection of *pCAG-ZZEF1-ALOX15-T2A-L2T* with pCMV-Cre in HEK293T cells produced strong tdTomato reporter fluorescence after 24 hours, confirming construct functionality in the presence of Cre (Figure 1B). Mice generated using this construct (*ZZEF1-ALOX15* mice) were crossed to *HRGP-Cre* and *Dkk3-Cre* driver lines, designed to activate transgene expression in cone photoreceptors or most retinal progenitor cells, respectively.

However, neither transgene activation nor a retinal phenotype was observed in *HRGP-Cre^+/-^ ;ZZEF1-ALOX15^+/-^* or *Dkk3-Cre^+/-^;ZZEF1-ALOX15^+/-^* mice. Retinal harvests from *HRGP-Cre^+/-^ ;ZZEF1-ALOX15^+/-^* and *Dkk3-Cre^+/-^;ZZEF1-ALOX15^+/-^* mice demonstrated successful Cre-lox recombination, evidenced by the excision of the LSL cassette located between the CMV early enhancer/chicken β-actin (CAG) promoter and the *ZZEF1-ALOX15* chimera (Figure 1C).

However, neither *HRGP-Cre^+/-^;ZZEF1-ALOX15^+/-^*nor *Dkk3-Cre^+/-^;ZZEF1-ALOX15^+/-^* mice exhibited outer retinal damage or lipofuscin deposition relative to controls on fundus and OCT examination, expected indicators of retinal damage (Figure 1D-F). While one (1 of 4) transgenic mouse (*HRGP-Cre^+/-^;ZZEF1-ALOX15^+/-^*) showed mild age-related drusenoid lesions at week 54, similar findings were observed in a control littermate (*HRGP-Cre^+/-^;ZZEF1-ALOX15^-/-^*). Live imaging with a tdTomato excitation/emission filter at week 5 did not detect tdTomato fluorescence in *Dkk3-Cre^+/-^;ZZEF1-ALOX15^+/-^* mice (Figure 1E). Also, immunohistochemistry of retinal cryosections at P11 (*Dkk3-Cre^+/-^;ZZEF1-ALOX15^+/-^*) and 8 weeks (*HRGP-Cre^+/-^ ;ZZEF1-ALOX15^+/-^*) similarly did not detect tdTomato signal or any morphological abnormalities (Figure 1F). In line with this, end-point RT-PCR failed to detect the *ZZEF1-ALOX15* fusion junction transcript from retinal cDNA of either Cre-activated line, despite detection of Cre transcript (Supplementary Figure S1). Collectively, the absence of tdTomato reporter activation, retinal pathology and RNA expression in *HRGP-Cre^+/-^;ZZEF1-ALOX15^+/-^* and *Dkk3-Cre^+/-^ ;ZZEF1-ALOX15^+/-^* mice is consistent with transgene silencing.

### Long-Read Sequencing Reveals Unintended Off-target Integration and Concatenation of the Transgene in the *ZZEF1-ALOX15* Mouse

While genotyping targeting the tdTomato reporter of founder mice and F1 carriers confirmed germline transmission of the transgene cassette (Supplementary Table S2), 3′ junction PCR (spanning the ∼4.3 kb *ROSA26* homology arm and into the endogenous locus) was consistently challenging to obtain and interpret, leaving uncertainty regarding the transgene’s integration site and structural architecture. To resolve this, we sequenced the genome of a *ZZEF1-ALOX15^+/-^* mouse, obtaining 428 GB of data consisting of 7.5 million HiFi reads. Alignment to the GRCm39 reference genome achieved a mean genome-wide coverage of 49*×* (>99.99% of primary HiFi reads mapped, corresponding to 7,502,968 of 7,503,051 reads, including 11,225 low-quality alignments). Furthermore, we identified 410 vector-derived reads from raw sequencing data upon alignment to vector component sequences (including 2 reads identified from the barcode-unassigned HiFi read dataset), corresponding to 0.005% of all reads generated.

Initial visualization of alignments using IGV revealed a disproportionately high depth of reads containing plasmid elements at the intended *ROSA26* locus, inconsistent with single-copy integration (Supplementary Figure S2A). These reads mapped to this locus solely through the two *ROSA26* homology arms present in the plasmid. Terminal sequences of these reads did not span into the endogenous locus, and alignment upon the removal of transgene-derived reads identified through our baiting strategy was consistent with diploid sequencing coverage at the locus (50× across *ROSA26* and 48× across the intended integration site within the locus) (Supplementary Figure S2B). This pattern suggested that construct integration had likely occurred elsewhere in the genome. Bait extraction and annotation of individual reads containing plasmid elements identified extensive plasmid-plasmid junctions, consistent with concatenation (Figure 2).

**Figure 2.**
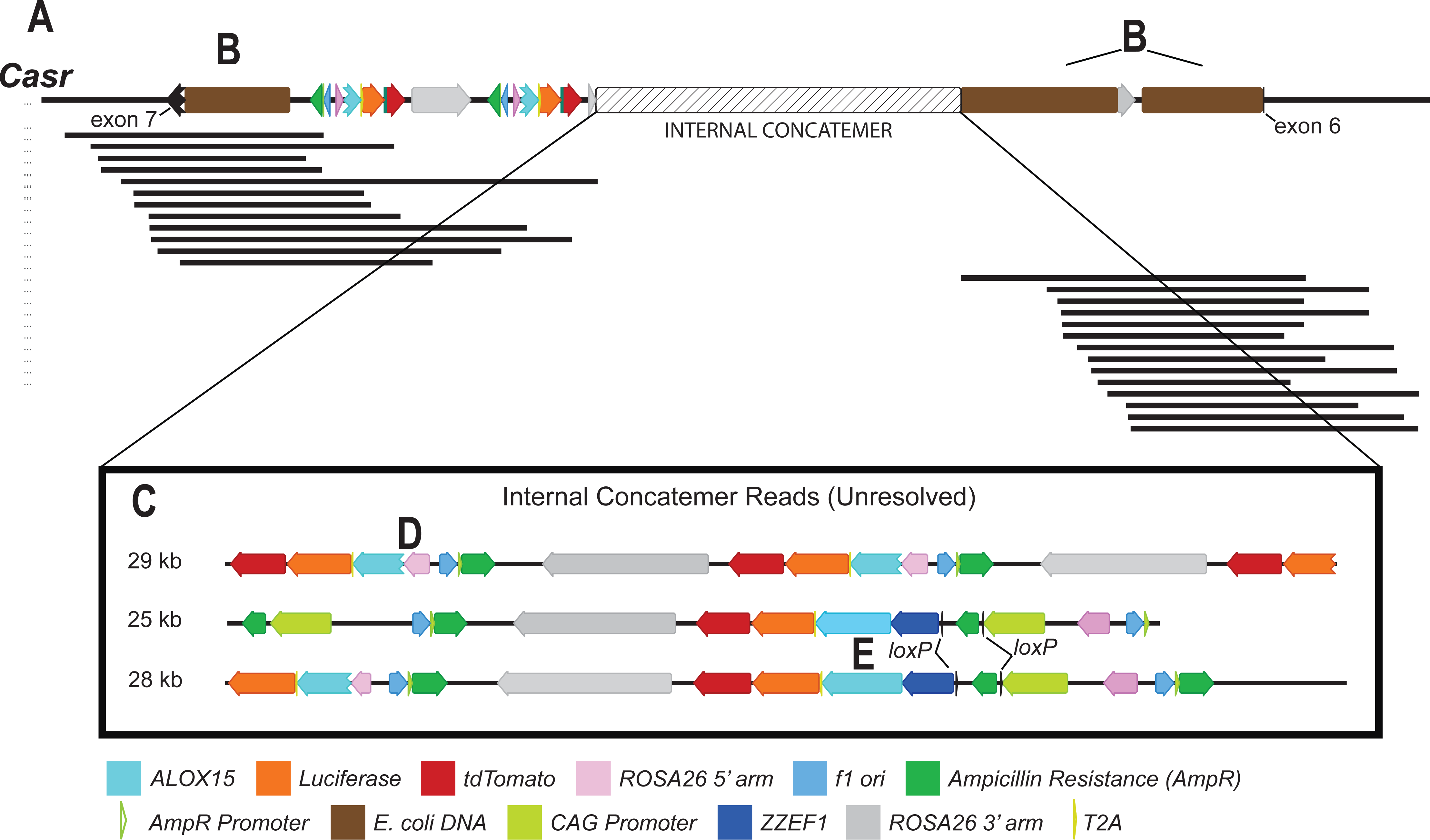
**(A)** Annotated consensus sequence generated using Geneious Prime from junction-spanning HiFi reads shows concatemer integration into *Casr*, accompanied by **(B)** *E. coli* genomic DNA at both breakpoints (7.7 kb at the 3′ junction and 8.8 kb at the 5′ junction). At the 5′ end of the concatemer, the 8.8 kb *E. coli* fragment is followed by a truncated 3′ *ROSA26* homology-arm segment and an additional 11.6 kb *E. coli* fragment. A 1.8 kb deletion spans exons 6 and 7 of *Casr*. **(C)** Individual reads sampled from within the concatemer (not aligned) demonstrate at least three head-to-tail transgene-derived copies and the presence of at least one structurally preserved expression cassette. The longest read supporting the concatemer was ∼29 kb. **(D, representative)** Multiple copies of the transgene (including at the 3′ junction) contain truncated *ZZEF1-ALOX15* fusion sequences and lack the CAG promoter. **(E)** At least one copy containing the structurally preserved expression cassette was identified.

Construct-derived reads with termini that did not match against a database of construct features were considered potential genome-transgene junctions and used to identify the concatemer integration site. We identified 26 junctional reads aligning to *Casr* on chromosome 16 (6.3% of all transgene-derived reads). Of these, 14 aligned to the 5′ junction and 12 to the 3′ junction of the concatemer (Figure 2A). The longest reads obtained from the 5′ and 3′ junction were 25 and 35 kb, respectively. Additional transgene integration sites were not detected among the vector-derived reads. IGV visualization of the *Casr* locus supported a heterozygous transgene integration which was coincident with a 1.8 kb deletion (Chr16:36,316,176-36,317,973) spanning from the terminus of exon 6 towards the start of exon 7 (Supplementary Figure S3). All reads corresponding to the integrated allele supported the 1.8 kb deletion, suggesting the deletion occurred concurrently with transgene integration. Furthermore, sequence homologies or microhomologies between the host-genome and exogenous sequences were not identified at either breakpoint nor were templated insertions or local rearrangements. Whereas the expression vector featured an LSL cassette and the *ZZEF1-ALOX15* chimera immediately downstream of the CAG promoter (Figure 1A), the two head-to-tail copies captured at the 3′ integration junction displayed substantial structural disruption, evidenced by the absence of the CAG promoter and *ZZEF1* sequences (Figure 2D). Furthermore, backbone vector elements such as the ampicillin resistance gene and a truncated *ROSA26* 5′ homology arm were discovered immediately upstream of the fusion gene, characterized by the complete loss of the *ZZEF1* sequence and partial truncation of *ALOX15*. Along with tdTomato and luciferase, this truncated fusion gene was oriented opposite to the direction of *Casr* transcription. *In silico* prediction using Cas-OFFinder did not identify any off-target Cas9 editing sites at the deletion site or elsewhere within the *Casr* locus.

Genome-wide SV calling with pbsv identified 28,647 quality-filtered variants (Supplementary Table S3). Among these, a ∼288 kb duplication on chromosome 9 (chr9:46,782,333-47,070,700) was found to overlap with a site predicted by Cas-OFFinder. However, given that the site contained four mismatched bases relative to the sgRNA and was located ∼53 kb from the nearest breakpoint, this duplication was not considered a likely Cas9-mediated off-target event. No other predicted off-target sites were found near an SV.

From within the internal concatemer, the longest read obtained (29 kb) captured three head-to-tail oriented copies of the transgene with structural losses like those observed at the integration junction, further highlighting the extent of the rearrangement beyond the expected 11 kb integration cassette (Figure 2C). Among all transgene-derived reads, we identified at least one copy of the structurally preserved expression cassette (Figure 1A), which was recovered from the internal concatemer rather than the junctional reads (Figure 2E). *De novo* assembly of the complete concatemer could not produce a plausible consensus sequence beyond the integration junctions, likely because the repeat array could not be contained within the read lengths obtained. Consequently, the orientation and precise location of the preserved copy within the overall structure remains unknown.

To explore the mechanism underlying transgene silencing from the structurally preserved cassette, we used pb-CpG-tools to perform 5-methylcytosine (5mC) profiling across CAG promoter sequences from ear-notch DNA. Using a modification-score cutoff of ≥50, 95.7% (176/184) of CpG positions across the CAG promoter were classified as methylated, consistent with promoter hypermethylation (mean modification score, 91.4; mean coverage, 43×; Figure 3). Specifically, 100% (20/20) of CpG positions within the CMV enhancer, 78% (25/32) in the chicken β-actin promoter, and 99.2% (131/132) in the chimeric intron were predicted to be methylated (Figure 3). Given that methylation status cannot be resolved in a copy-specific manner, this profiling represents the aggregate methylation state across all CAG elements present. Given a mean genome-wide coverage of 49×, the observed coverage at the CAG sequence is consistent with approximately two copies of the promoter present within the concatemer. We note that 12/52 CpG sites located across the CMV enhancer and chicken β-actin promoter produced modification scores between 36-66 (Data File S4). This may reflect differing methylation status across copies at these sites or local CpG-specific variability within copies.

**Figure 3.**
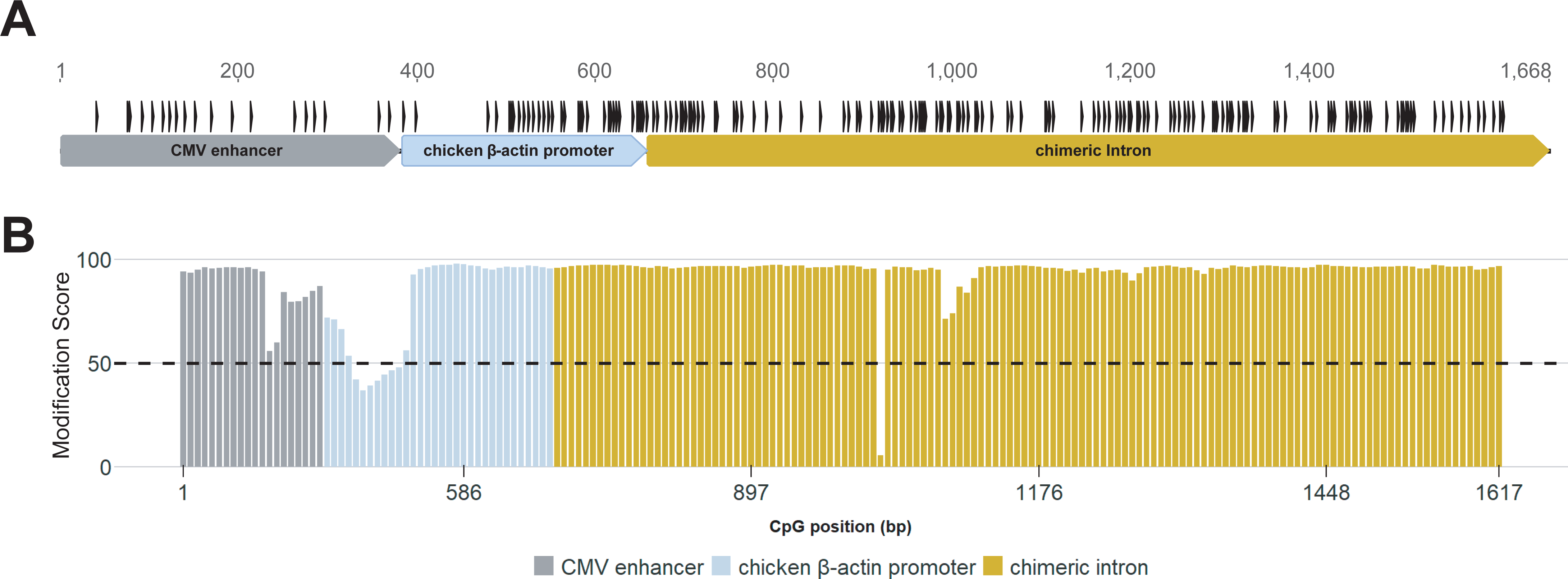
**(A)** Schematic showing three regions of the CAG promoter in pCAG-ZZEF1-ALOX15-T2A-L2T, generated using Geneious Prime. CpG dinucleotides are indicated by black vertical bars. **(B)** Bar chart showing the aggregate 5mC modification scores (1-100) of CpG dinucleotides across all copies of the CAG promoter present within the concatemer, indicating hypermethylation within copies of the transgene promoter. Methylation status was called from PacBio HiFi reads using Jasmine and parsed using pb-CpG-tools. Colored bars denote sites within the CMV enhancer, chicken β-actin promoter, and chimeric intron regions. The horizontal dashed line indicates a modification score threshold of 50 determining CpG dinucleotide methylation status.

Nevertheless, for 87% (160/184) of CpG sites across the promoter, ≥90% of supporting reads were concordant in methylation status, suggesting a high degree of consistency between copies of the CAG promoter overall. Thus, our data indicates widespread hypermethylation across CAG promoter sequences within the concatemer. The annotated output file generated using pb-CpG-tools, reporting modification score and coverage for each CpG site, has been provided in Data File S4.

Notably, BLAST analysis of concatemer-flanking sequences revealed co-integration with *E. coli* genomic DNA contaminants (Figure 2B). At the 3′ integration junction, we identified a 7.7 kb fragment containing 5 neighboring genes from the *E. coli* genome (*pgl, ybhD, ybhH, ybhI* and *ybhJ)* (Supplementary Figure S4). A second 8.8 kb fragment was detected at the 5′ junction, comprising 6 genes (*ybhA, modA, modB, modC, galE* and *galT).* Notably, these two segments map to a single contiguous region of the *E. coli* DH5α genome (Supplementary Figure S5).

Downstream of the fragment at the 5′ junction, we additionally observed a partial 1.3 kb remnant of the 3′ *ROSA26* homology arm followed by a distinct 11.6 kb *E. coli* DNA fragment comprising 11 genes (*acrF, acrE, acrS, yhdU, yhdJ, fis, dusB, prmA, panF, yhdT,* and *accC*) (Supplementary Figure S4). Thus, we identified vector concatemerization and unintended heterozygous off-target integration in *Casr* on chromosome 16, with rearrangements involving contaminating *E. coli* DNA contributing to final transgene architecture.

### Aberrant Construct Integration at *Casr* is Associated with Altered Calcium Homeostasis

Breakpoint PCR of the 5′ and 3′ junctions confirmed the heterozygous transgene integration across exons 6 and 7 of *Casr* in *ZZEF1-ALOX15* mice used for biochemical and expression analysis (Supplementary Figure S6). Exploratory qPCR analysis demonstrated an approximately 77% reduction of retinal *Casr* expression in a *ZZEF1-ALOX15^+/-^* mouse relative to a sex-matched wild-type control (*ZZEF1-ALOX15^-/-^*; Supplementary Figure S7). CaSR senses changes to extracellular calcium (Ca^2+^) concentration and plays a central role in maintaining systemic homeostasis, partly through parathyroid hormone (PTH) secretion^46^. Heterozygous *Casr* knockout (KO) mice recapitulate familial hypocalciuric hypercalcemia (FHH), a clinically benign condition characterized by modest elevations in serum [Ca^2+^] and reduced urinary [Ca^2+^] excretion^47,48^. To assess the physiological consequences of the genomic disruption at the *Casr* locus, we measured serum and urine [Ca^2+^] in a cohort of *ZZEF1-ALOX15* mice. We observed ∼15% higher serum [Ca^2+^] in these mice relative to control littermates (11.17 ± 0.87 mg/dL vs. 9.70 ± 1.23 mg/dL) (Figure 4A). After adjusting for age and sex, transgenic genotype was associated with a 1.25 mg/dL increase in serum calcium (95% CI, 0.18-2.32 mg/dL; p = 0.025). Urinary calcium excretion (normalized to creatinine) in our model (0.26 ± 0.15) was not significantly altered in comparison to control littermates (0.19 ± 0.03) after adjustment for age and sex (β = 0.023; 95% CI, −0.10 to 0.15; *p* = 0.70) (Figure 4B). This finding was unchanged in a sensitivity analysis excluding the single high-value calcium-to-creatinine ratio observed in a *ZZEF1-ALOX15* mouse (β = 0.021; 95% CI, −0.06-0.10; p = 0.57). The age, sex, and raw calcium measurements of all animals used for biochemical analysis are provided in Data File S5.

**Figure 4.**
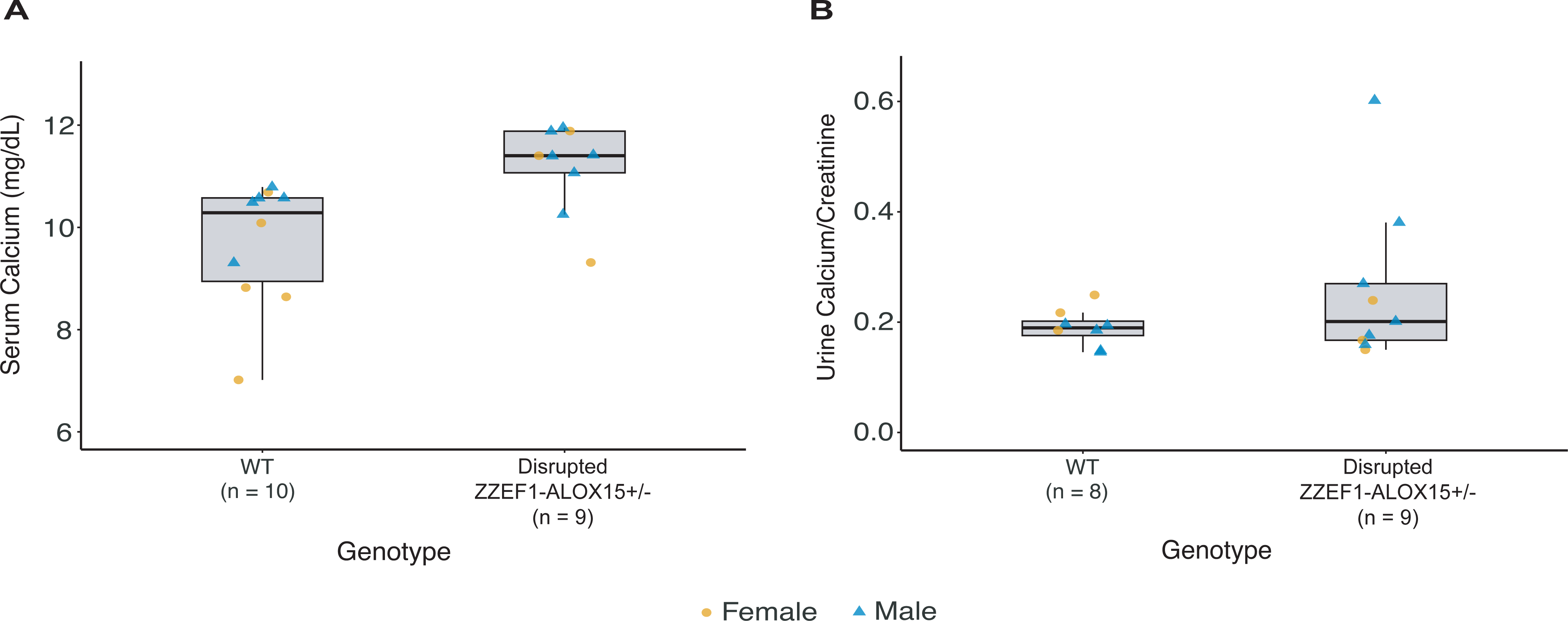
**(A)** Box plots of serum calcium concentration (mg/dL) in littermate controls lacking the transgene (n = 10) and disrupted *ZZEF1-ALOX15+/-* mice (n = 9). The disrupted line exhibited higher serum calcium levels compared with controls after adjusting for age and sex (multiple linear regression; β = 1.25 mg/dL; 95% CI, 0.18-2.32 mg/dL; *p* = 0.025). **(B)** Box plots of urinary calcium to creatinine ratios (Ca/Cr) in littermate controls (n = 8) and disrupted *ZZEF1-ALOX15+/-* mice (n = 9). Urinary Ca/Cr in the disrupted line did not differ significantly in comparison to control mice after adjusting for age and sex (multiple linear regression; β = 0.023; 95% CI, −0.10 to 0.15; *p* = 0.70). Horizontal lines within each box indicate median values, 25th and 75th percentiles are denoted by upper and lower fences, and whiskers extend to 1.5× the interquartile range. Individual dots denote calcium measurements for each mouse and are color- and shape-coded by sex.

## Discussion

We identified multi-copy concatenation and unintended off-target integration of the transgene into the *Casr* locus using whole-genome LRS. Consequently, aggregate 5mC profiling from ear-notch DNA indicated hypermethylation across copies of the CAG promoter within the concatemer, providing a potential epigenetic contribution for transgene silencing observed in the retina of *ZZEF1-ALOX15* mice from functional assays. Finally, biochemical analysis indicated that these rearrangements caused an unintended phenotype in the model partially consistent with heterozygous *Casr* knockout mice^47,49^.

Indeed, we observed no evidence of *ZZEF1-ALOX15* chimera expression at the RNA level or tdTomato reporter fluorescence in the disrupted *ZZEF1-ALOX15* model, despite evidence of *in vitro* transgene activation from the same construct. Mouse retina previously electroporated with *ZZEF1-ALOX15* under the same promoter context (*pCAG-ZZEF1-ALOX15-IRES-GFP*) demonstrated rosette formation and thinning of the outer nuclear layer by 8 weeks, consistent with severe retinal damage^32^. In contrast, retinal lamination in these disrupted *ZZEF1-ALOX15* mice appeared unremarkable in comparison to their littermate controls, inconsistent with the expected HMD phenotype. Together, these findings implicate transgene concatenation and off-target integration into *Casr* behind transgene silencing of the germline model.

Although the molecular basis of concatenation remains poorly understood, transgene concatemers primarily assemble through homologous recombination^50,51^. Host DNA repair mechanisms can join identical transgene copies following cell microinjection leading to tandemly ligated and truncated expression cassettes prior to genomic integration^10^. As most reads analyzed similarly exhibited shortened transgene copies and lacked critical regulatory elements such as the CAG promoter, expression failure could be explained by structural disruption.

However, the presence of at least one structurally preserved expression cassette suggests alternative silencing mechanisms. To this end, repeat-induced gene silencing (RIGS) is a well-documented consequence of concatenation and is thought to occur through repressive epigenetic modifications^10^. Repetitive elements are highly regulated in mammalian cells and are typically correlated with CpG methylation^52,53^. In the context of transgenic modelling, CAG promoter-driven expression is known to be negatively correlated with methylation of CpG dinucleotides within the CMV enhancer and chicken β-actin intron^54,55^. Accordingly, CAG promoter copies within the concatemer were predominantly methylated across these regions. We therefore hypothesize that transgene silencing in the model was multifactorial, likely attributable to RIGS and truncations within the concatemer leading to the loss of the CAG promoter within some copies of the transgene.

We performed biochemical analysis in disrupted *ZZEF1-ALOX15* mice to assess the phenotypic consequences of *Casr* disruption. *Casr* encodes a dimeric G-protein-coupled receptor that is highly expressed within the parathyroid gland and the kidneys^46,56,57^. Within the parathyroid gland, the receptor exerts its calciotropic effect by repressing parathyroid hormone (PTH) in response to elevated serum [Ca^2+^], thereby inhibiting bone resorption and stimulating renal Ca^2+^ excretion^56^. Evidence suggests that local CaSR modulation in the kidneys occurs independently of PTH and involves downregulation of claudins 16 and 19^49^. Mice carrying heterozygous loss-of-function variants in *Casr* recapitulate features of familial hypocalciuric hypercalcemia, characterized by [Ca^2+^] that is modestly elevated in the blood but low in urinary excretion^47^. We found altered serum calcium homeostasis in *ZZEF1-ALOX15* disrupted mice, suggesting that transgene integration compromised CaSR function. Interestingly, despite elevated serum calcium, urinary calcium levels were not significantly different in these mice relative to controls. The phenotype observed in our disrupted model closely resembles the recently reported *Casr*^BCH002^ model, which carries an inactivating variant within the transmembrane domain of *Casr* and similarly exhibits elevated serum calcium with inappropriately normal urinary calcium excretion^49^. Although the mechanisms underlying this intermediate phenotype remain unknown, our exploratory analysis in our model demonstrated reduced *Casr* expression at the RNA level. We therefore speculate that heterozygous *Casr* disruption in *ZZEF1-ALOX15* mice may alter calcium homeostasis and contribute to an intermediate phenotype through haploinsufficiency.

Suggestive of double-strand break (DSB) repair outcomes, we identified a 1.8 kb genomic deletion at the integration site spanning from the terminus of exon 6 to the start of exon 7 in *Casr*. Junctional architecture at the integration breakpoints, devoid of homology/microhomology, local rearrangements and templated insertions, was most consistent with non-homologous end joining (NHEJ). Given that *in silico* analysis using Cas-OFFinder did not predict Cas9 editing off-target sites in the *Casr* locus, the underlying source of the break remains unknown.

Reconstruction of junctional reads revealed the incorporation of bacterial DNA at both breakpoints. Such co-integrations have been previously reported and it is likely these rearrangements occurred with contaminating *E. coli* DNA present during model generation^26^. Notably, analysis of 40 widely used mouse lines revealed that 25% (10/40) harbored *E. coli* DNA alongside their transgenes^7^. Given that only 5% of transgenic strains published on the Mouse Genome Database have mapped integration sites, such rearrangements are considered to be more widespread than presently recognized^7^.

While our study underscores the value of unbiased whole-genome LRS in characterizing complex transgene rearrangements, certain limitations warrant consideration. Although Cre-lox recombination demonstrating excision of the loxP-flanked stop cassette between the CAG promoter and *ZZEF1* was detected in both Cre-activated transgenic lines (Figure 1C), whether recombination occurred within a structurally preserved expression cassette could not be conclusively ascertained. Furthermore, while plausible off-target effects were not identified among the sites predicted by Cas-OFFinder, the possibility that some of the remaining SVs resulted from Cas9-mediated editing cannot be fully excluded due to the absence of a matched control dataset. Additionally, as CAG confers ubiquitous activity across most tissue types^58,59^, the extensive hypermethylation observed across the promoter from ear-notch DNA was aberrant and strongly suggestive of transgene silencing. Nevertheless, the methylation profile obtained from this tissue may not fully reflect that of the retina. Finally, given that *ZZEF1-ALOX15^+/-^* mice were only rederived by our group, the original founder was not available for retrospective testing. Therefore, circumstances underlying the original quality-control assessment at MICAM, such as potential founder mosaicism harboring both a *ROSA26* allele and off-target transgene concatemer, are incompletely understood.

Several strategies for evaluating transgenic mouse models have been proposed, including conventional PCR and targeted locus amplification (TLA)^13,28^. However, these methods provide limited resolution when substantial deviations from transgene structure occur and may require supplementary testing, such as confirmatory sequencing or methylation analysis, to inform model viability^28^. While our analysis combined unbiased LRS with targeted transcript analysis to evaluate transgene viability, integrating whole-genome analysis with transcriptomic or proteomic profiling may provide an alternative approach for assessing the downstream consequences of transgene rearrangements during model troubleshooting or validation^8,60–62^. In our case, challenges in reconciling the technical limitations of junctional PCR with erroneous integration were evident, as routine quality control failed to detect the unintended rearrangement. As shown here and by others, LRS can facilitate comprehensive and efficient transgene integration analysis^15,63–65^. Indeed, our approach generated sufficient read and methylation data to resolve both junction sites and guide model troubleshooting, though complete reconstruction of the concatemer was not possible, as read lengths did not span the full repeat array. The choice of LRS-based mapping strategy remains largely dependent on cost, efficiency, and experiment objective, with Cas9 enrichment paired with ONT LRS currently presenting a lower-cost option with simplified downstream analysis^28,66^. Nonetheless, as recent developments in LRS technology, such as the Revio system from PacBio, continue to improve sequencing throughput while reducing per-genome costs, unbiased LRS examination is likely to become increasingly accessible, thereby enabling routine global interrogation of edited genomes. To this end, our data demonstrates the growing feasibility of unbiased whole-genome LRS as a powerful tool for the quality control of transgenic mouse models.

## Declaration statements

### 1. Data Availability

The raw sequencing data generated in this study were deposited in the NCBI Sequence Read Archive (SRA) under BioProject accession number PRJNA1471078 and are available at the following URL: https://www.ncbi.nlm.nih.gov/bioproject/PRJNA1471078.

## 2. Acknowledgements

We extend our gratitude to the staff at The Centre for Phenogenomics, including the Pathology Core for their assistance with serum calcium measurements. MM was awarded a scholarship from the Vision Science Research Program (Department of Ophthalmology and Vision Sciences, University of Toronto). P.P.M. holds the Anne and Max Tanenbaum Chair in Neuroscience (Krembil Research Institute) and E.H. holds the Henry Brent Chair in Innovative Pediatric Ophthalmology (The Hospital for Sick Children, Toronto).

## 4. Author Contributions

**M.M.:** Methodology, Investigation, Visualization, Writing - Original Draft, Writing - Review and Editing. **K.A.:** Methodology, Validation, Supervision, Writing - Review and Editing. **R.H.:** Investigation, Visualization. **E.T.:** Methodology, Validation, Supervision, Writing - Review and Editing. **Z.B.:** Investigation, Writing - Review and Editing. **R.A.:** Investigation, Writing - Review and Editing. **A.D.:** Investigation, Writing - Review and Editing. **B.G.:** Resources, Writing - Review and Editing. **M.D.W.:** Methodology, Writing - Review and Editing. **E.I.:** Methodology, Writing - Review and Editing. **P.P.M.**: Methodology, Resources, Writing - Review and Editing. **E.H.:** Methodology, Writing - Review and Editing. **A.V.:** Conceptualization, Methodology, Validation, Supervision, Writing - Review and Editing, Funding Acquisition.

