## Supplementary Figures and Tables for "Unbiased Long-Read Whole-Genome Sequencing Enables High-Resolution Mapping of Transgene Concatenation and Off-target Genomic Disruption in a Mouse Model"

### Supplementary Table S1. CRISPR-Cas9 Components Used for Model Generation

| Component | Name | Sequence (5' – 3') |
|---|---|---|
| **gRNA** | ***ROSA26* gRNA** | ACTCCAGTCTTTCTAGAAGA |

### Supplementary Table S2. List of Oligos Organized by Experiment

| **Oligo ID** | **Sequence 5' – 3'** |
|---|---|
| **Founder/F1 Quality Control (*ROSA26* Targeting Validation)** | |
| To identify WT (320 bp) from mutant (600 bp) mice | |
| R26 A | AAG ACC GCG AAG AGT TTG TC |
| R26 B | AAA GTC GCT CTG AGT TGT TAT |
| R26 C | GGA GCG GGA GAA ATG GAT ATG |
| For 3' Junction (4483 bp) | |
| Vin V2 | GGT TTT TTA ATT CGC GGC CCT AGA AG |
| R26 R2 | CTG AGA CCA TTC TCA GTG GCT CAA |
| For 3' Wild-type Allele (4547 bp) | |
| TF | CTC TGC TGC CTC CTG GCT TCT |
| R26 R2 | CTG AGA CCA TTC TCA GTG GCT CAA |
| For 5' Junction (1325 bp) | |
| Rosa26 F4 | CTT GGT GCG TTT GCG GGG AT |
| SR62 R | CCA CTG GAA AGA CCG CGA AGA GTT |
| For 5' Wild-type Allele (1338 bp) | |
| Rosa26 F4 | CTT GGT GCG TTT GCG GGG AT |
| TF | CTC TGC TGC CTC CTG GCT TCT |
| **Germline Transmission (PCR)** | |
| Genotyping of tdTomato Reporter (734 bp) | |
| P5785 (F) | GGC GAG GAG GTC ATC AAA GA |
| P5786 (R) | TCC GAG GAC AAC AAC ATG GC |
| **Verification of Cre-lox Recombination (PCR)** | |
| P6370 (F) | CTC CAT CTC CAG CCT CGG |
| P6371 (R) | ATG CCT CGA CAC CAG CG |
| **Genotyping of Cre-driver lines (PCR)** | |
| For HRGP-Cre (412 bp) | |
| P6599 (F) | AGG TGT AGA GAA GGC ACT TAG C |
| P6600 (R) | CTA ATC GCC ATC TTC CAG CAG G |
| For Dkk3-Cre (685 bp) | |
| P6487 (F) | CAG ACC ATA CTA GTT TGG CAG TAC TGG GA |
| P6488 (R) | CTT GCG AAC CTC ATC ACT CGT TGC A |
| **Determination of Transgene and Cre Expression (PCR)** | |
| For the *ZZEF1-ALOX15* fusion junction (146 bp) | |
| P6457 (F) | GGT GAC GGC TTC TAT GGA GA |

| | |
|---|---|
| P6458 (R) | CTA TGC CGG TTC CAA CAA CC |
| For Cre (412 bp) | |
| P6599 (F) | AGG TGT AGA GAA GGC ACT TAG C |
| P6600 (R) | CTA ATC GCC ATC TTC CAG CAG G |
| **To Identify 5' and 3' Transgene-*Casr* Breakpoints (PCR)** | |
| For 5' Wild-type Allele (410 bp) | |
| 5' WT (F) | AGG GAC ACA CTA TTG GAC GG |
| 5' WT (R) | TCC CTT GAT GCC CCA ATT CTA |
| For 5' Junction (477 bp) | |
| 5' TG (F) | GTG ATG GCA GCC GAT TAT GAA A |
| 5' WT (R) | TCC CTT GAT GCC CCA ATT CTA |
| For 3' Junction (501 bp) and Wild-type Allele (295 bp) | |
| 3' WT (F) | TGG TGG GTA TTT TGG CCT CA |
| 3' WT (R) | GCT CTC ACT CTC TTT GCG GT |
| 3' TG (R) | AGA GCG CGA GAT TAT CAA GG |
| ***Casr* Expression Analysis by qPCR** | |
| For *Casr* exons 2-3 (166 bp) | |
| *Casr* F | GTC GTC GGC TGC AAT TGT G |
| *Casr* R | TAT CCC CCA GGT GAG CTA CG |
| For normalization to *Ppib* (88 bp) | |
| *Ppib* F | ACG AGT CGT CTT TGG ACT CTT T |
| *Ppib* R | GCC AAA TCC TTT CTC TCC TGT A |

**Supplementary Table S3. Summary of Structural Variants Discovered by pbsv**

| SV Type | All pbsv Calls | Quality-filtered (PASS) | Quality-filtered SVs ≥10 kb |
|---|---|---|---|
| **Insertion** | -- | 13,950 | 25 |
| **Deletion** | -- | 11,230 | 25 |
| **Duplication** | -- | 3,320 | 7 |
| **Inversion** | -- | 37 | 10 |
| **Breakend** | -- | 100 | NA |
| **Copy-number Variant** | -- | 10 | 9 |
| **Total** | 29,007 | 28,647 | 76 |

A

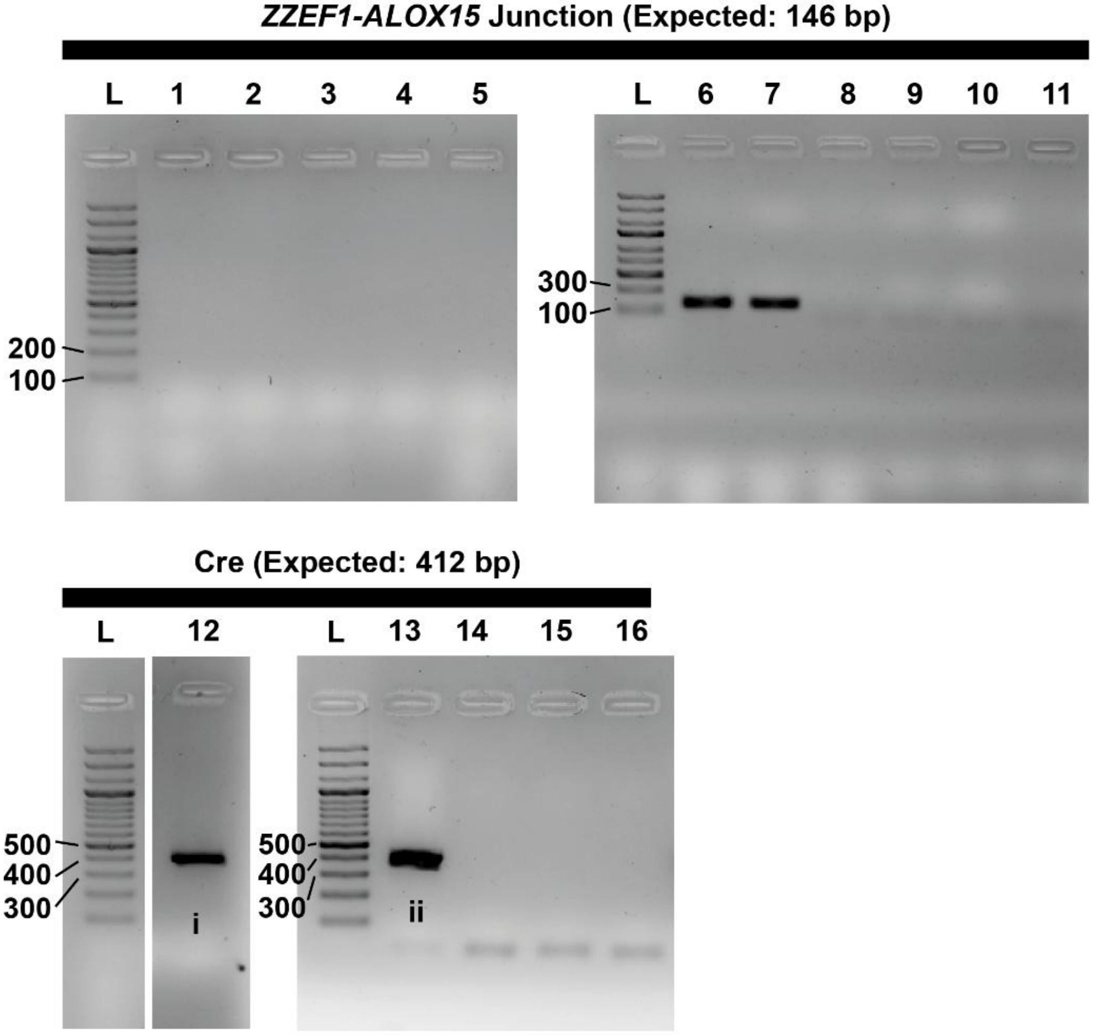


1. *HRGP-Cre+/-;ZZEF1-ALOX15+/-* (retinal cDNA)
2. *Dkk3-Cre+/-;ZZEF1-ALOX15+/-* (retinal cDNA)
3. No RT control (*Dkk3-Cre+/-;ZZEF1-ALOX15+/-*)
4. No template control
5. NTC
6. Positive technical control
7. Positive technical control
8. Non-transfected negative control
9. No RT control (*Dkk3-Cre+/-;ZZEF1-ALOX15+/-*)
10. No template control
11. NTC
12. *HRGP-Cre+/-;ZZEF1-ALOX15+/-* (retinal cDNA)
13. *Dkk3-Cre+/-;ZZEF1-ALOX15+/-* (retinal cDNA)
14. No RT control (*Dkk3-Cre+/-;ZZEF1-ALOX15+/-*)
15. No template control
16. NTC

B

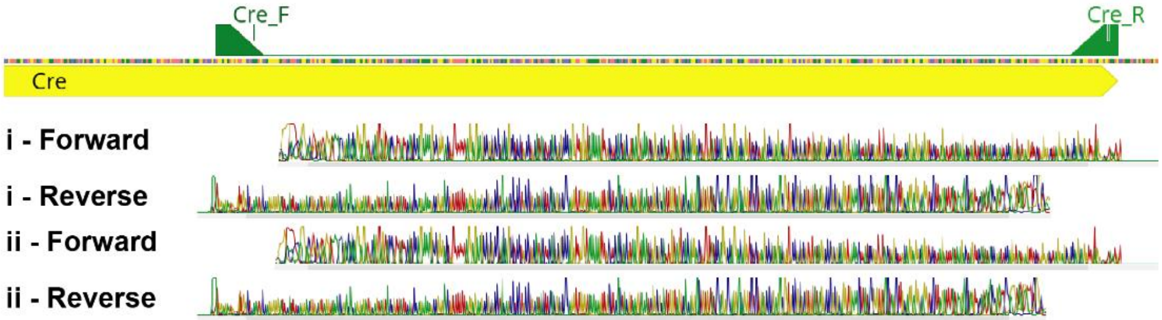

**Supplementary Figure S1.** Transgene expression analysis from mice carrying both Cre and the *ZZEF1-ALOX15* transgene. **(A)** Gel electrophoresis following PCR amplification from retinal cDNA shows no observable amplification of the *ZZEF1-ALOX15* fusion junction from *HRGP-Cre*$^{+/-}$*;ZZEF1-ALOX15*$^{+/-}$ and *Dkk3-Cre*$^{+/-}$*;ZZEF1-ALOX15*$^{+/-}$ mice. A product corresponding to Cre was observed in both mice. Positive technical controls were obtained using cDNA from HEK293T cells co-transfected with a construct carrying *ZZEF1-ALOX15* under control of the CAG promoter. **(B)** Sanger sequencing alignment (Geneious Prime) confirming the identity of the gel-purified Cre amplicons from both mice.

A

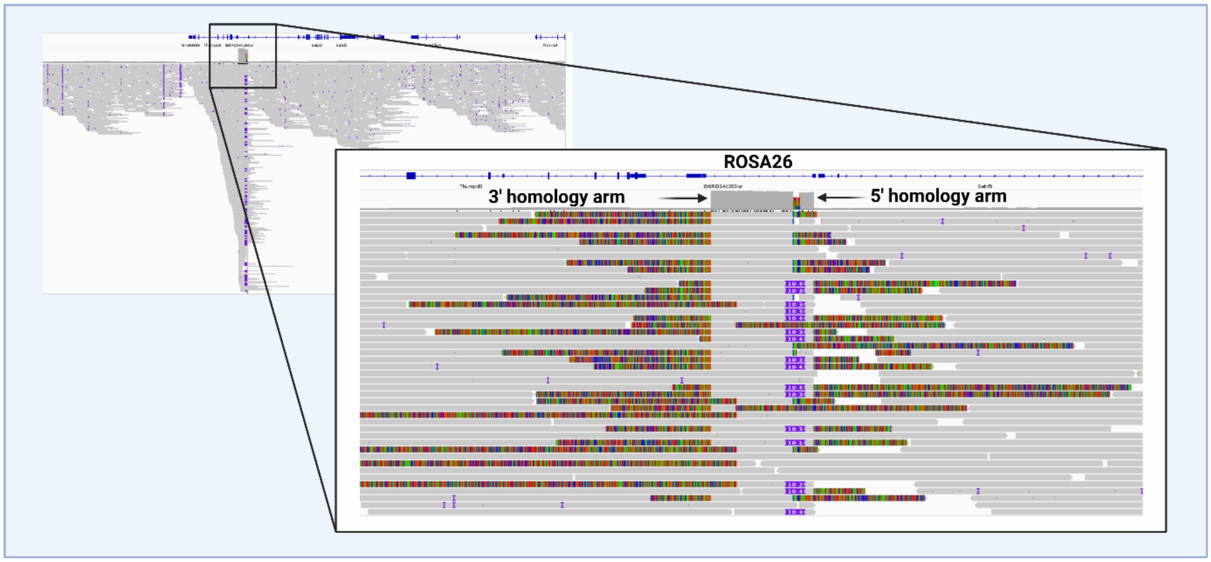


B

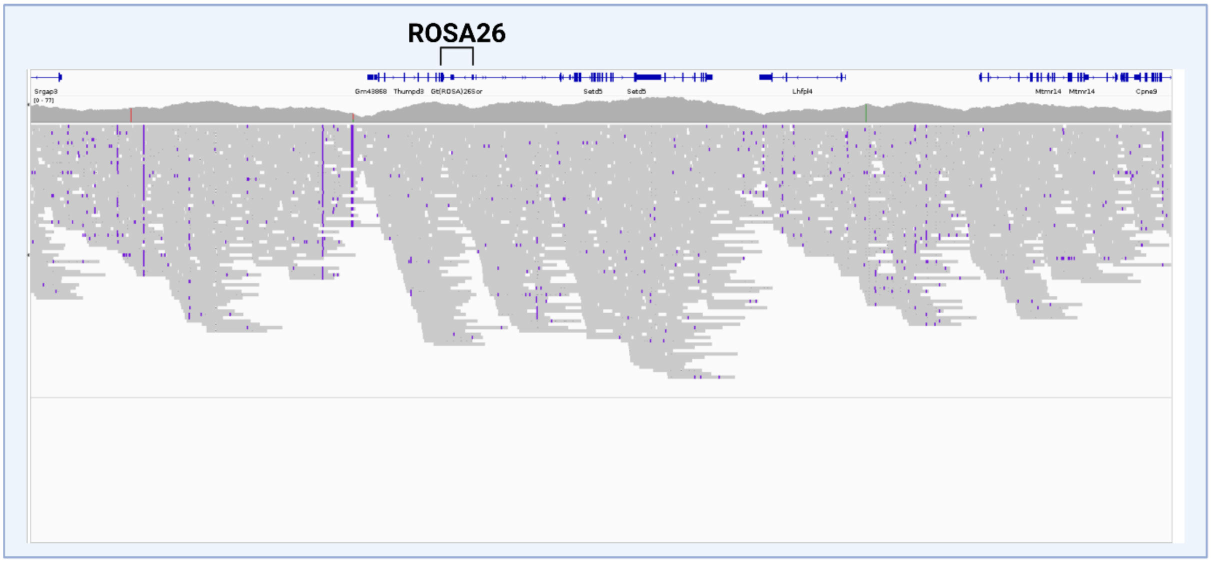


**Supplementary Figure S2. (A)** IGV visualization of HiFi reads at *ROSA26* locus on chromosome 11. BLAT results of mismatched reads identified plasmid-derived sequences. Arrows on coverage depth track indicate plasmid containing reads mapping to regions corresponding to the right and left homology arms. **(B)** IGV visualization of the *ROSA26* locus upon removal of plasmid-derived reads. The resulting depth of coverage at the locus was consistent with the average genome diploid sequencing coverage.

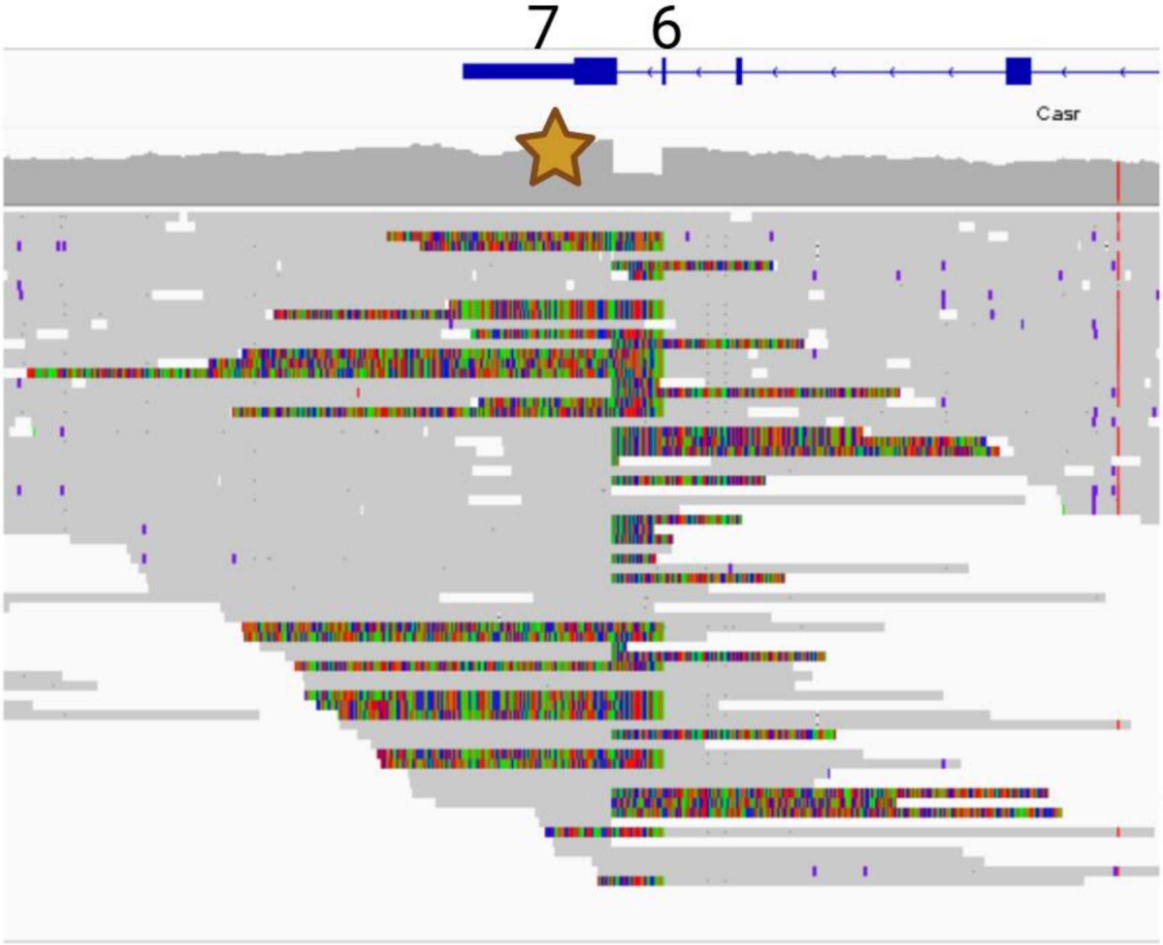


**Supplementary Figure S3.** IGV visualization of the *Casr* locus on chromosome 16. A sharp drop in read depth (★) is observed between exons 6 and 7 (Chr16:36,316,176-36,317,973), corresponding to a ~1.8 kb deletion. BLAT results of mismatched reads show plasmid-derived sequences which span into the *Casr* locus. Consistent with heterozygous transgene integration, the 5′ and 3′ integration junctions were supported by 21 and 25 reads respectively, while 25 reads were found to span the corresponding WT allele. All reads corresponding to the transgene allele supported the 1.8 kb deletion, suggesting that the deletion coincided with the transgene integration. Read depth across the integration was higher than that obtained across the breakpoints through our baiting strategy (14 and 12 at the 5' and 3' breakpoints, respectively) as multiple reads at the locus strictly overlapped with *Casr* and the flanking *E. coli* sequences without extending into the transgene and hence were not included in the baiting strategy.

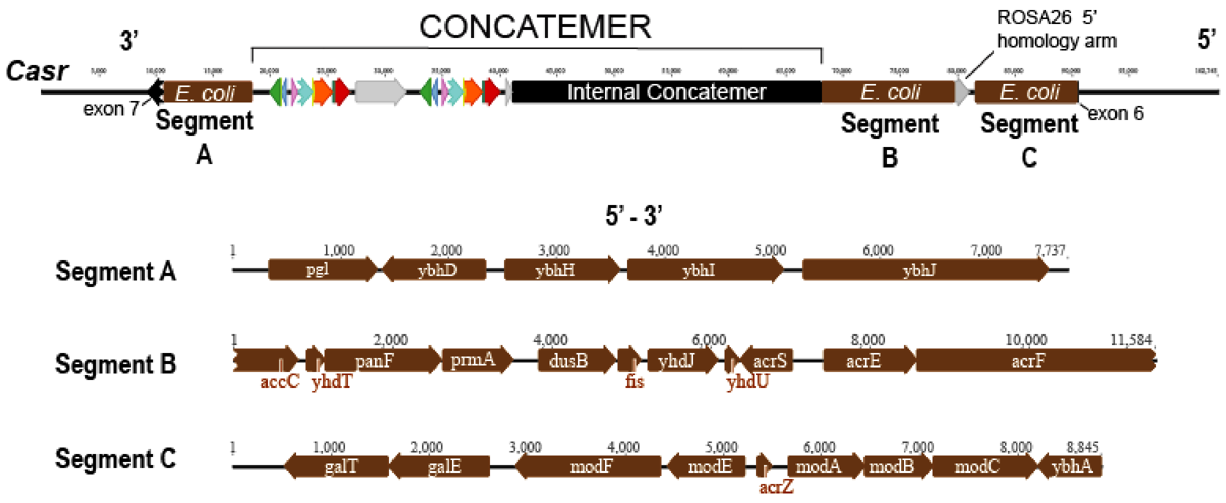


**Supplementary Figure S4.** Schematic of three *E. coli* genomic fragments (A, B, and C) that co-integrated with the concatenated transgene at the *Casr* locus. Corresponding annotations show the *E. coli* coding sequences contained within each fragment.

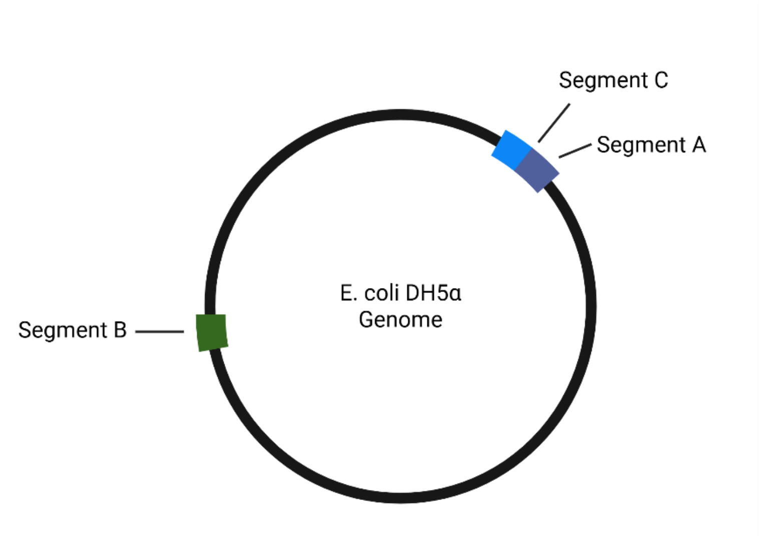


**Supplementary Figure S5.** Representation of the DH5α genome showing the relative positions of the three co-integrating fragments (A, B, and C). Fragments C and A originate from a contiguous region of the genome.

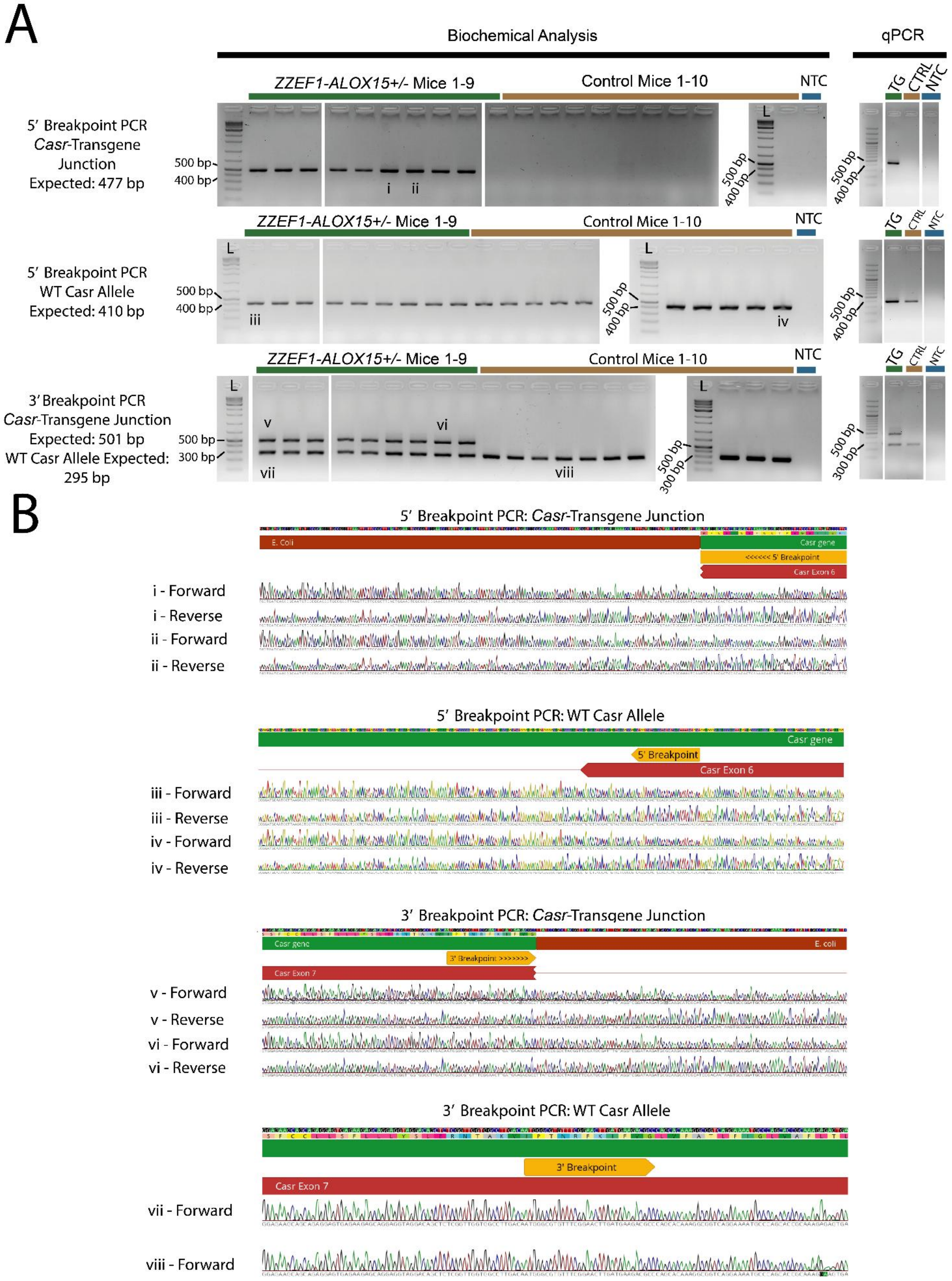
A
Biochemical Analysis
qPCR
ZZEF1-ALOX15+/- Mice 1-9
Control Mice 1-10
NTC
TG
CTRL
L
5′ Breakpoint PCR
Casr-Transgene
Junction
Expected: 477 bp
500 bp
400 bp
i
ii
5′ Breakpoint PCR
WT Casr Allele
Expected: 410 bp
iii
iv
3′ Breakpoint PCR
Casr-Transgene Junction
Expected: 501 bp
WT Casr Allele Expected:
295 bp
300 bp
v
vi
vii
viii
B
5′ Breakpoint PCR: Casr-Transgene Junction
E. Coli
Casr gene
<<<<<< 5′ Breakpoint
Casr Exon 6
i - Forward
i - Reverse
ii - Forward
ii - Reverse
5′ Breakpoint PCR: WT Casr Allele
5′ Breakpoint
iii - Forward
iii - Reverse
iv - Forward
iv - Reverse
3′ Breakpoint PCR: Casr-Transgene Junction
E. coli
3′ Breakpoint >>>>>>>
Casr Exon 7
v - Forward
v - Reverse
vi - Forward
vi - Reverse
3′ Breakpoint PCR: WT Casr Allele
3′ Breakpoint
vii - Forward
viii - Forward

*L, DNA ladder; TG, transgenic mouse (ZZEF1-ALOX15$^{+/-}$); CTRL, control mouse (ZZEF1-ALOX15$^{-/-}$); NTC, no-template control.*

**Supplementary Figure S6.** PCR and Sanger sequencing validation of the transgene-*Casr* integration in the mice used for biochemical analysis (n = 9 *ZZEF1-ALOX15*$^{+/-}$, n = 10 control *ZZEF1-ALOX15*$^{-/-}$) and exploratory qPCR analysis of *Casr* expression (n = 1 *ZZEF1-ALOX15*$^{+/-}$, n = 1 control *ZZEF1-ALOX15*$^{-/-}$). **(A)** Agarose gel electrophoresis of PCR targeting the 5′ and 3′ breakpoints of the transgene integration within the *Casr* locus. Detection of the disrupted allele and corresponding wild-type product in *ZZEF1-ALOX15* mice demonstrates heterozygous transgene integration within the *Casr* locus. **(B)** Sanger sequencing of select PCR products confirming the 5′ and 3′ breakpoints in the disrupted (i-ii, v-vi) and homologous wild-type *Casr* allele (iii-iv, vii-viii). The yellow labelled "breakpoint" region indicates the genomic position corresponding to the integration breakpoint observed in the disrupted allele. Gel electrophoresis images were generated by the ChemiDoc Imaging system (Bio-Rad, Hercules, CA, USA). Sanger chromatograms were visualized in Geneious Prime.

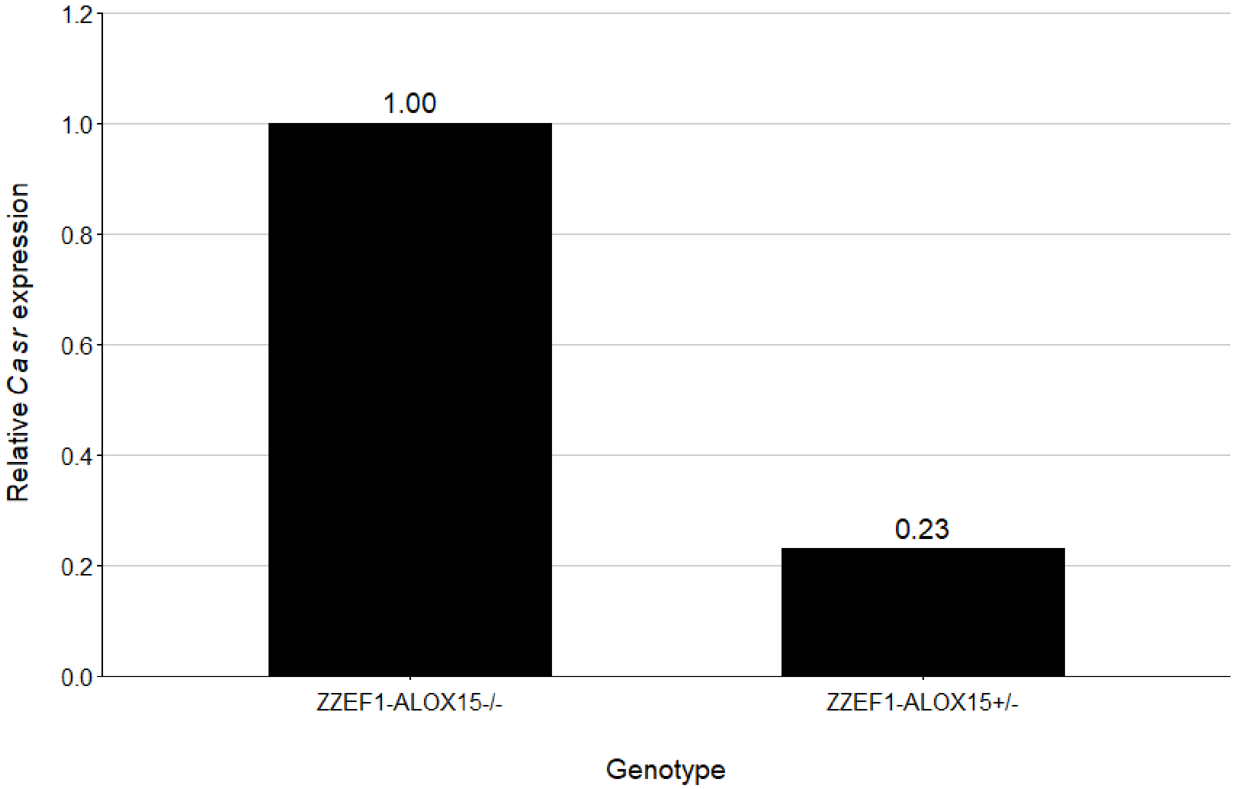


**Supplementary Figure S7.** Exploratory qPCR analysis from retinal cDNA showing reduced *Casr* expression in a Dkk3-*Cre*$^{-/-}$;*ZZEF1-ALOX15*$^{+/-}$ mouse relative to a sex-matched *ZZEF1-ALOX15*$^{-/-}$ control. *Casr* expression was normalized to *Ppib*, with WT expression set to 1. One biological sample was run in technical triplicates for each genotype.

**Supplementary Data File Legends**

**Data File S1.** Construct map of *pCAG-ZZEF1-ALOX15-T2A-L2T*, provided as a GenBank file.

**Data File S2.** A multi-FASTA of the 12 sequence baits used to identify transgene-derived sequences.

**Data File S3.** Key workflows, tools, and annotated commands used for HiFi read alignment, baiting, and CpG methylation analysis.

**Data File S4.** Site-level CAG promoter methylation output generated by pb-CpG-tools, with annotated CAG promoter regions.

**Data File S5.** Sex, age, genotype, and biochemical data for all animals included in serum calcium and urinary calcium-to-creatinine ratio analyses.
