## Supplementary material for "Unbiased Long-Read Whole-Genome Sequencing Enables High-Resolution Mapping of Transgene Concatenation and Off-target Genomic Disruption in a Mouse Model": Data Files S1-S5: Data File S3.pdf

Workflows for HiFi read alignment, baiting, and CpG methylation analysis are provided below. All commands were executed from a Bash shell using a high-performance computing cluster configured with SAMtools (version 1.21), minimap2 (version 2.24), pbmm2 (version 26.1.0), pbsv (version 2.6.2) and pb-CpG-tools (version 3.0.0). Geneious Prime (version 2025.1.3) was used for the baiting and methylation analysis workflows.

#### Section 1. Alignment of PacBio HiFi Reads to the Mouse Reference Genome

1. Index the GRCm39 mouse (*Mus musculus*) genome assembly for minimap2 alignment.

```
# Reference Genome: Mus musculus GRCm39 genomic FASTA (NCBI RefSeq
accession GCF_000001635.27)

minimap2 -d Mus_musculus.GRCm39.dna.primary_assembly.fa.mmi
GCF_000001635.27_GRCm39_genomic.fna
```

2. Convert the raw BAM file to FASTQ format for minimap2 alignment.

```
# Convert HiFi BAM to FASTQ for minimap2 alignment

samtools fastq m84090_250425_202437_s2.hifi_reads.bc2072.bam >
HifiReads.fastq
```

3. Compress the FASTQ file using gzip.

```
# Compression was performed to reduce storage requirements

gzip HifiReads.fastq
```

4. Align HiFi reads to the GRCm39 reference genome and sort records by genomic coordinates.

```
# Minimap2 alignment using the 'map-pb' preset, with soft clipping
enabled (-Y)

minimap2 -t 16 -ax map-pb -Y
Mus_musculus.GRCm39.dna.primary_assembly.fa.mmi HifiReads.fastq.gz
| samtools sort -@ 4 -m 6G -o aligned_HifiReads.bam
```

5. Create a BAM index file to enable fast lookup of alignments by genomic position.

```
samtools index aligned_HifiReads.bam
```

### Section 2. Genome-wide Structural Variant Calling

1. Identify structural variants from the “aligned\_HifiReads.bam” file generated in step 1.4.

```
#discover structural variation signatures

pbsv discover aligned_HifiReads.bam HifiReads_discover.svsig.gz

#call structural variants

pbsv call Mus_musculus.GRCm39.dna.primary_assembly.fa
HifiReads_discover.svsig.gz structural_variants.vcf
```

2. Retain variants marked as “FILTER=PASS” in the resulting “structural\_variants.vcf” file.
3. Compare the genomic coordinates of retained pbsv calls with the off-target sites predicted by Cas-OFFinder.

### Section 3. Read Baiting and Transgene Breakpoint Reconstruction

1. Partition the *pCAG-ZZEF1-ALOX15-T2A-L2T* sequence into 12 non-overlapping bins spanning the construct using the “Extract Region” function in Geneious Prime. Export the baits as a single multi-FASTA file (Plasmid\_Baits.fasta, provided in Data File S2). Sequences solely corresponding to the *ROSA26* homology arms on the construct were excluded as standalone baits to prevent capturing solely endogenous reads lacking other construct elements.

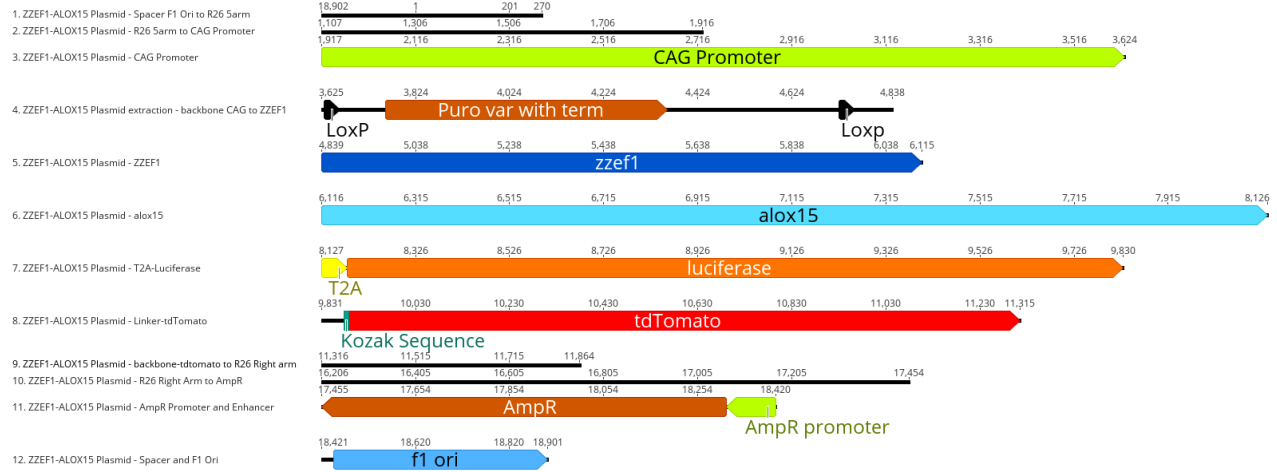

- Align the gzip-compressed FASTQ file generated from step 1.3 to the sequence bait multi-FASTA reference.

```
# Minimap2 alignment using the 'map-pb' preset, with soft clipping
enabled (-Y) and secondary alignments suppressed (--secondary=no).

minimap2 -t 16 -ax map-pb -Y --secondary=no Plasmid_Baits.fasta
HifiReads.fastq.gz | samtools view -h -F 4 - | samtools sort -@ 4 -m
6G -o baited_reads.bam
```

- Generate a “Read to Bait” table to view all unique bait associations (read-bait pairs).

```
samtools view baited_reads.bam | awk '{print $1"\t"$3}' | sort |
uniq > read_to_bait_table.txt
```

- Remove reads corresponding to endogenous murine orthologues of *ZZEF1* or *ALOX15*.

```
# This step removes reads solely identified by the ZZEF1 or ALOX15
baits (ZZEF1-ALOX15_Plasmid_-_ZZEF1 and ZZEF1-ALOX15_Plasmid_-
_alox15, respectively). Reads captured by either ZZEF1 or ALOX15
baits in addition to at least one other transgene bait are retained.

grep -v -E "ZZE1-ALOX15_Plasmid_-_alox15|ZZE1-ALOX15_Plasmid_-_
_ZZE1" read_to_bait_table.txt > read_to_bait_table_noaloxzze1.txt
```

- To account for the edge case where a transgene-derived read was captured by both the *ZZE1* and *ALOX15* baits but by no other transgene bait, reads shared by the *ZZE1* and *ALOX15*

baits were identified and added to `read_to_bait_table_noaloxzzef1.txt` prior to deduplication.

```
# Generate a list of all reads captured by the ALOX15 bait

awk '$2=="ZZEF1-ALOX15_Plasmid_-_alox15"{print $1}'
read_to_bait_table.txt > alox15_reads.txt

# Generate an equivalent list of all reads captured by the ZZEF1
bait

awk '$2=="ZZEF1-ALOX15_Plasmid_-_ZZEF1"{print $1}'
read_to_bait_table.txt > zzef1_reads.txt

# Find reads present in both lists

grep -Ff alox15_reads.txt zzef1_reads.txt > shared_alox15_zzef1.txt

# Add reads shared by both ZZEF1 and ALOX15 baits to the table
generated in 3.4 which removed reads solely generated by the ZZEF1
or ALOX15 baits

awk '{print $1"\tshared_alox15_zzef1"}' shared_alox15_zzef1.txt >>
read_to_bait_table_noaloxzzef1.txt

# Generate final deduplicated list of baited reads

cut -f1 read_to_bait_table_noaloxzzef1.txt | sort | uniq >
final_trusted_reads.txt
```

6. Extract transgene-derived reads from the baited BAM file generated in step 3.2 using the list of deduplicated baited reads generated in step 3.5.

```
# Use final deduplicated list of reads

samtools view -N final_trusted_reads.txt -b baited_reads.bam >
baited_reads_cleaned.bam

# Convert the resulting BAM file into FASTQ format before importing
into Geneious Prime

samtools fastq baited_reads_cleaned.bam > baited_reads_cleaned.fastq
```

9. Create a subfolder within the “Reference Features” directory containing the 12 sequence baits generated in step 3.1.
10. Annotate reads in “`baited_reads_cleaned.fastq`” using the “Annotate & Predict” tool, applying a 50% sequence similarity threshold against the subfolder.
  • Sequence similarity thresholds ranging from 30-90% were tested. For this particular dataset, a 50% cutoff was found to best capture truncated or corrupted transgene-associated elements while controlling for sequence misidentification. Depending on the application and research context, this threshold may require custom optimization.
11. Perform BLAT queries of baited reads with unresolved terminal sequences to identify junction-spanning reads and transgene integration site(s).
  • Sequences were manually extracted and queried against the GRCm39 *Mus musculus* reference genome using BLAT. Candidate genomic integration sites were identified based on the longest, highest-identity genomic alignments. *Casr* was consistently identified across all reads with unresolved terminal sequences. No other genomic loci were detected among the transgene-derived reads. Identification of all 5' and 3' junction reads was performed using “Annotate & Predict”, annotating reads against *Casr* with a minimum sequence similarity of 50%. This threshold was used for feature identification and not as the initial standard for integration site discovery. Any remaining unresolved internal sequences were further queried using BLAST.
12. Repeat steps 3.2-3.11 for the raw barcode-unassigned HiFi read dataset.
  • Two additional transgene-derived reads were identified from this file and appended to the final list of transgene-derived reads.
13. Upon finding no evidence of transgene integration at the *ROSA26* locus (given the absence of *ROSA26*-transgene junction-spanning reads), inspect baited reads within Geneious Prime to remove the *ROSA26*-derived sequences with no additional construct-specific sequences.
  • These reads were attributed to the WT *ROSA26* locus captured by the “Spacer F1 Ori to R26 5arm” bait seen in step 3.1 which contains an overhanging segment of the *ROSA26* 5' homology arm.
14. Export a FASTQ file containing the finalized 410 transgene-derived reads as “`transgene_reads.fastq`” for inspection of the *ROSA26* locus (steps 4.1-4.3).
15. Separate breakpoint-spanning HiFi reads into 5' and 3' junction groups for independent *de novo* assembly using the built-in Geneious assembler (medium sensitivity/fast settings). Afterwards, generate a consensus sequence of the resulting contig using the “Generate Consensus Sequence” tool, applying a “Highest Quality (60%)” threshold.

16. Align the 5' and 3' junction consensus sequences independently to the *Casr* reference. Identify each integration breakpoint as the boundary between the portion of the consensus sequence aligned to *Casr* and the soft-clipped sequence corresponding to exogenous DNA.
17. Append the soft-clipped exogenous sequence from each junction consensus to the corresponding *Casr* breakpoint to generate an extended integration-site reference. Remap the junction-spanning HiFi reads to the reconstructed reference to generate and validate the final integration-site map.

##### Section 4. Removal of Transgene-Derived Reads from Aligned BAM File for Inspection of the *ROSA26* locus

1. Obtain a list of read IDs from the FASTQ file generated in step 3.14.

```
# Generate a list of read IDs by extracting FASTQ read headers and
remove the leading '@' from each ID

grep '^@' transgene_reads.fastq | sed 's/^@//' >
transgene_read_ids.txt
```

2. Remove transgene-derived reads from the alignment and then index the resulting BAM file using SAMtools.

```
# Select reads (-N) whose IDs are listed in transgene_read_ids.txt
and extract those not selected (-U) into
aligned_HifiReads.no_transgene.bam

samtools view -b -U aligned_HifiReads.no_transgene.bam -N
transgene_read_ids.txt aligned_HifiReads.bam > /dev/null

# Generate a BAM index file

samtools index aligned_HifiReads.no_transgene.bam
```

3. Import “aligned\_HifiReads.no\_transgene.bam” into IGV to verify that coverage at the *ROSA26* locus is consistent with no transgene integration.

### Section 5. 5mC Profiling of the Transgene CAG Promoter

1. Generate a deduplicated list of read IDs captured by the CAG promoter bait.

```
# -u removes duplicate IDs
awk '$2=="ZZEF1-ALOX15_Plasmid_-_CAG_Promoter" {print $1}'
read_to_bait_table.txt | sort -u > cag_read_ids.txt
```

2. Extract CAG promoter-containing reads from the raw barcode-assigned HiFi read dataset into a separate BAM file.

```
samtools view -@ 8 -b -N cag_read_ids.txt
m84090_250425_202437_s2.hifi_reads.bc2072.bam -o cag_reads.bam
```

3. Highlight the CAG promoter from the construct in Geneious Prime and export it as a FASTA file to serve as an alignment reference.
4. Index the custom CAG promoter reference sequence using pbmm2.

```
pbmm2 index CAG_promoter.fa CAG_promoter.mmi
```

5. Align the BAM file containing unique CAG promoter reads to the indexed reference.

```
pbmm2 align CAG_promoter.mmi cag_reads.bam cag_reads.CAGonly.bam --
preset CCS --sort
```

6. Remove secondary and supplementary alignments.

```
# Remove (-F) records containing secondary or supplementary
alignment flags (0x900)

samtools view -b -F 0x900 cag_reads.CAGonly.bam -o
aligned.primary_nosupp.bam
```

7. Index the BAM file.

```
samtools index aligned.primary_nosupp.bam
```

8. Process the filtered BAM file using pb-CpG-tools.

```
#Outputs compressed BED and BigWig files for analysis of CpG
methylation data

aligned_bam_to_cpg_scores --bam aligned.primary_nosupp.bam --output-
prefix CAG_methylation --threads 12
```

9. Retrieve per-site modification scores from the resulting BED file.
10. Annotate CpG dinucleotides across the CAG promoter reference using Geneious Prime (Annotate & Predict > Find Motifs) and export the position of each site using the “Export Table” function under the annotations tab.
11. Cross-check CpG dinucleotides identified by pb-CpG-tools against those identified by Geneious Prime.
12. Classify CpG sites with modification scores $\geq 50$ as methylated.
